# Total number and ratio of GABAergic neuron types in the mouse lateral and basal amygdala

**DOI:** 10.1101/2021.03.15.435365

**Authors:** Viktória K. Vereczki, Kinga Müller, Éva Krizsán, Zoltán Máté, Zsuzsanna Fekete, Laura Rovira-Esteban, Judit M. Veres, Ferenc Erdélyi, Norbert Hájos

## Abstract

GABAergic neurons are key circuit elements in cortical networks. In spite of growing evidence showing that inhibitory cells play a critical role in the lateral (LA) and basal (BA) amygdala functions, neither the number of GABAergic neurons nor the ratio of their distinct types have been determined in these amygdalar nuclei. Using unbiased stereology, we found that the ratio of GABAergic neurons in the BA (22 %) is significantly higher than in the LA (16 %) in both male and female mice. No difference was observed between the right and left hemispheres in either sexes. In addition, we assessed the ratio of the major inhibitory cell types in both amygdalar nuclei. Using transgenic mice and a viral strategy for visualizing inhibitory cells combined with immunocytochemistry, we estimated that the following cell types together compose the vast majority of GABAergic cells in the LA and BA: axo-axonic cells (5.5-6 %), basket cells expressing parvalbumin (17-20 %) or cholecystokinin (7-9 %), dendrite-targeting inhibitory cells expressing somatostatin (10-16 %), NPY-containing neurogliaform cells (14-15 %), VIP and/or calretinin-expressing interneuron-selective interneurons (29-38 %) and GABAergic projection neurons expressing somatostatin and neuronal nitric oxide synthase (nNOS, 5.5-8 %). Our results show that these amygdalar nuclei contain all major GABAergic neuron types as found in other cortical regions. Furthermore, our data offer an essential reference for future studies aiming to reveal changes in GABAergic cell number and in inhibitory cell types typically observed under different pathological conditions, and to model functioning amygdalar networks in health and disease.

**Significance statement:** GABAergic cells in cortical structures, like in the lateral and basal nucleus of the amygdala, have a determinant role in controlling circuit operation. In this study, we provide the first estimate for the total number of inhibitory cells in these two amygdalar nuclei. In addition, our study is the first to define the ratio of the major GABAergic cell types present in these cortical networks. Taking into account that hyper-excitability in the amygdala, arising from the imbalance between excitation and inhibition typifies many altered brain functions including anxiety, posttraumatic stress disorder, schizophrenia and autism, uncovering the number and ratio of distinct amygdalar inhibitory cell types offers a solid base for comparing the changes in inhibition in pathological brain states.

## Introduction

The amygdala is a complex structure that is composed of several functionally distinct nuclei. One of the main amygdalar regions, the basolateral amygdaloid complex (BLA) plays a key role in a variety of behavioral functions, including goal-directed and social behavior, formation and storage of affective memory as well as controlling the direction of attention (Phelps and LeDoux, 2005; Phelps et al., 2014; Janak and Tye, 2015; Gothard, 2020). The BLA, which is a nuclear extension of the cortex deep in the temporal lobe, consists of the lateral (LA), basal (BA) and accessory basal nuclei (Pitkanen et al., 1997). As in other cortical regions, the BLA principal cells giving rise to both local axonal collaterals and projections to remote target areas use glutamate for fast neurotransmission (Smith and Pare, 1994; McDonald, 1996; Pare and Smith, 1998; Pitkanen et al., 2003). In addition to these excitatory cells there are neurons in the BLA that release GABA as the main neurotransmitter molecule (McDonald, 1985; McDonald and Augustine, 1993). Previous studies obtained in rat and monkey estimated the number of GABAergic cells to be around 15 % and 25 % of the total neuronal population, respectively (McDonald, 1992; McDonald and Augustine, 1993). However, in these studies only male animals were investigated at a given BLA level in one of the hemispheres without separating the distinct BLA nuclei. Thus, it is unexplored whether there are differences in the GABAergic cell number between the nuclei, sexes and/or hemispheres.

Cortical inhibitory neurons are remarkably heterogeneous in their morphological, molecular and functional features (Kepecs and Fishell, 2014; Pelkey et al., 2017; Huang and Paul, 2019). Previous studies have established several cardinal GABAergic cell types that can be ubiquitously identified in all cortical regions and can be characterized by typical neurochemical content or a combination of these markers (Kepecs and Fishell, 2014). Axo-axonic cells, either containing or lacking the calcium binding protein parvalbumin (PV), target specifically the axon initial segment of excitatory principal neurons (Somogyi, 1977; He et al., 2016). Two types of basket cells innervating predominantly the soma and spine-free proximal dendrites of neurons express either PV or a neuropeptide cholecystokinin (CCK) and cannabinoid receptor type 1 (CB1)(Freund and Katona, 2007). Dendrite-targeting interneurons can be also separated into two types: one, containing somatostatin (SST), forms synaptic contacts predominantly with the distal dendrites of neurons (Kawaguchi and Kubota, 1996), while the other one, the so-called neurogliaform cell, often expressing neuropeptide Y (NPY), is responsible for slow inhibition of dendrites (Tamas et al., 2003). Another group of GABAergic cells is formed by interneuron-selective interneurons innervating specifically, if not exclusively, other GABAergic neurons and may contain vasoactive intestinal polypeptide (VIP) and/or the calcium binding protein calretinin (CR)(Acsády et al., 1996; Gulyas et al., 1996). Finally, there are GABAergic neurons that, in addition to their local axonal collateralization, project to remote brain areas and often show immunoreactivity for SST and neuronal nitric oxide synthase (nNOS)(Gulyas et al., 2003; He et al., 2016). Both the number and the ratio of distinct GABAergic neuron types show substantial variability among cortical regions (Kim et al., 2017) that may have significant computational consequences (Harris and Shepherd, 2015). Therefore, these circuit parameters in amygdalar networks should be determined to better understand circuit operation in the BLA.

Our study has been conducted in the LA and BA, the two nuclei that play distinct roles in various amygdala functions (Janak and Tye, 2015; Manassero et al., 2018) and may differ in their inhibitory circuits (Polepalli et al., 2020). For unbiased stereology, we visualized the GABAergic neurons in the mouse brain by intercrossing vesicular GABA transporter (Vgat)-Cre transgenic mice with reporter mice. To determine the fractions of distinct types of GABAergic neurons, we combined labeling of genetically defined neuronal populations in transgenic mice with immunocytochemistry. These approaches allowed us to accurately estimate the number of GABAergic neurons and the ratio of their types in the LA and BA.

## Materials and Methods

### Animals

All procedures involving animals were performed according to methods approved by the Hungarian legislation (1998. XXVIII. section 243/1998, renewed in 40/2013) and institutional guidelines. All procedures were in compliance with the European convention for the protection of vertebrate animals used for experimental and other scientific purposes (ETS number 123). Every effort was taken to minimize animal suffering and the number of animals used. For this study, the following mouse lines were obtained from The Jackson Laboratory (Bar Harbor, ME, USA) or from MMRRC (Table 1). To study CCK-expressing GABAergic neurons, Vgat-IRES-Cre mice were bred with BAC-CCK-GFPcoIN_sb mice and the offspring (Vgat^Cre^;CCK-GFPcoIN) were used in experiments. Two lines of BAC-CCK-GFPcoIN_sb mice differing in the BAC transgene copy number were generated similarly as BAC-CCK-DsRedT3 mice (Mate et al., 2013). In the offspring of the one copy line intercrossed with Vgat-IRES-Cre, GFP expression had lower levels in comparison to those offspring generated by crossing the two copy line with Vgat-IRES-Cre mice. Electrophysiological measurements were obtained in the offspring of BAC-CCK-GFPcoIN_sb mice having one or two copies of BAC transgene, whereas the interneuron counting was performed only in offspring of the two copy mouse line.

**Table 1.**
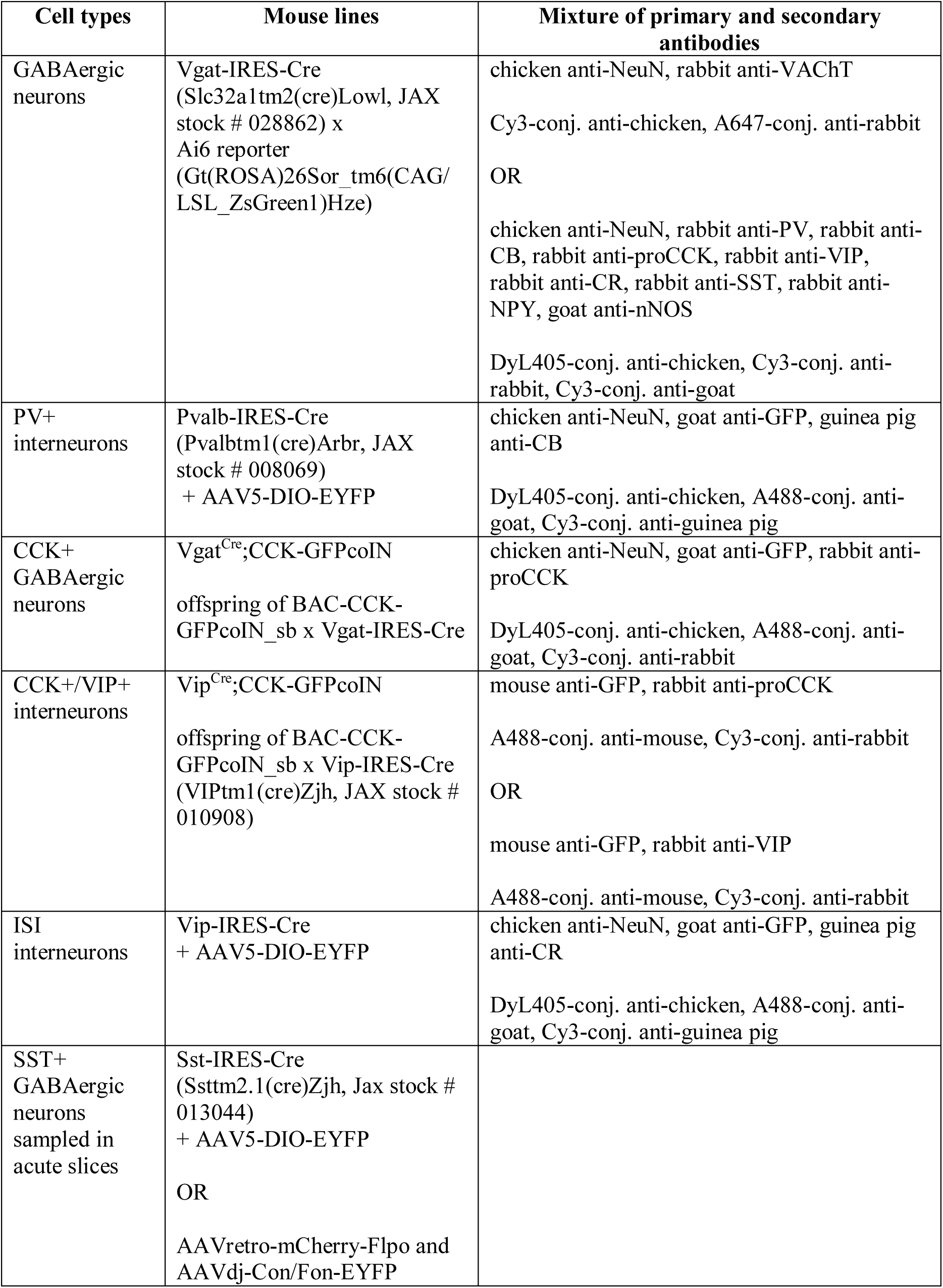
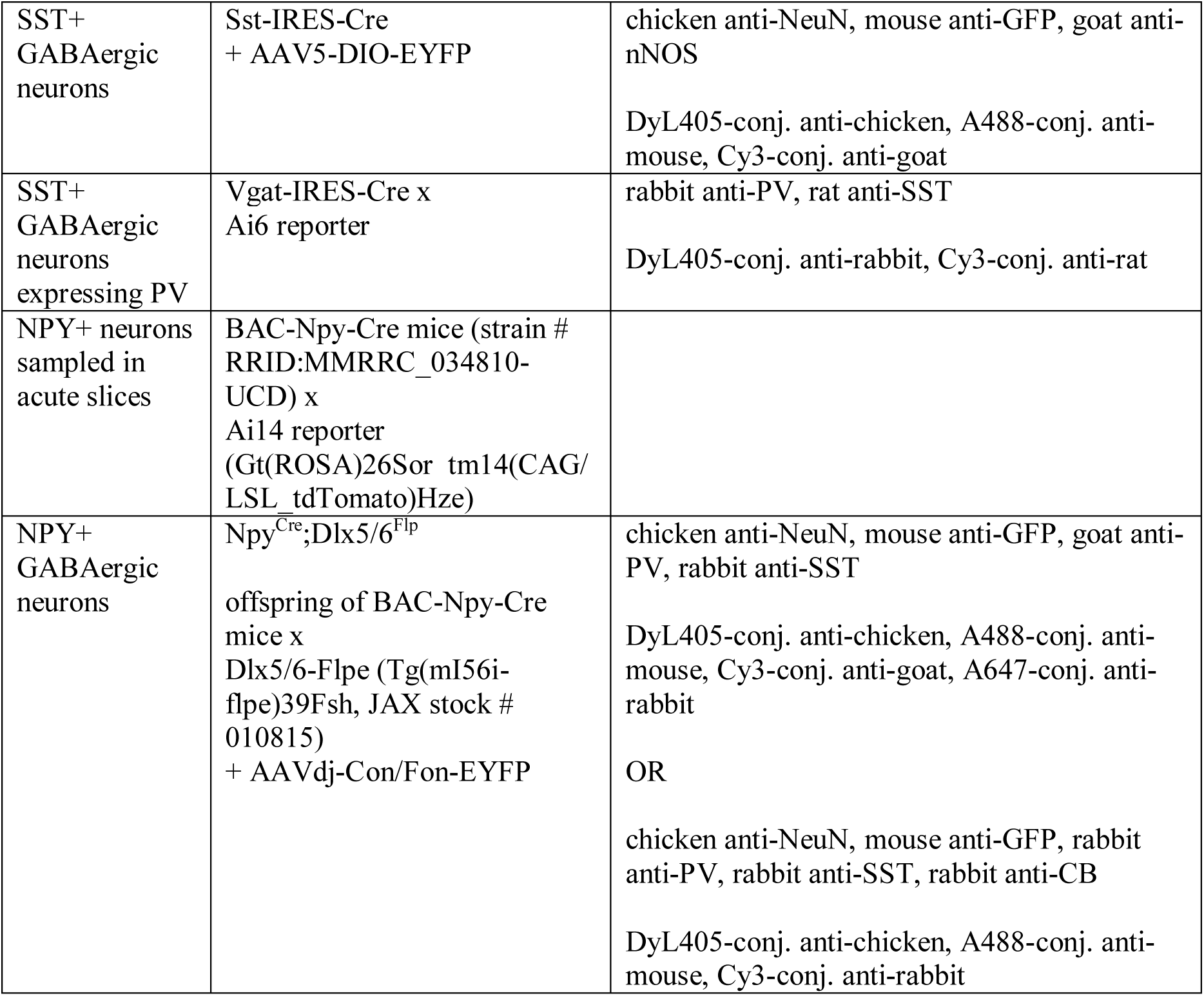
To estimate the total number of GABAergic neurons and the ratio of distinct inhibitory cell types in the LA and BA, the following mouse lines, AAVs and combinations of antibodies to visualize the given antigens were used in this study.

Males and females were used for stereology and electrophysiological recordings, while only the right hemisphere of the male mice were used to estimate the proportion of distinct interneuron types in each case. Mice were housed in same-sex groupings (2-4 per cage). Housing was in a temperature- and humidity-controlled *vivarium* under a 12 h light/dark cycle (lights on 06:00 h).

### Stereological Analysis

Brain tissue samples for stereological analyses were taken from 3 males and 3 females that were offspring of homozygous Vgat-IRES-Cre mice crossed with homozygous Ai6. After being anesthetized with ketamine/xylazine, adult mice (P56-70) were transcardially perfused with 0.9 % NaCl for 1-2 min followed by a fixative solution containing 4 % paraformaldehyde in 0.1 M phosphate buffer (PB; pH= 7.4) for 20 min. Coronal sections (50-µm thick) were prepared from the tissue blocks containing the entire amygdalar region using a Leica VT1000S vibratome (Leica Microsystems). Sections were stored in a cryoprotectant antifreeze solution consisting of glycerol, ethylene glycol, distilled H_2_O, and phosphate-buffer saline (3:3:3:1 volume ratio) at -20 °C until further processing (Watson et al., 1986). Using a random starting point within the amygdala, six or seven sections per animal containing the BLA were selected. The sections separated by 250 µm in rostro-caudal extent were immunostained for a neuronal marker, NeuN and for vesicular acetylcholine transporter, VAChT (Table 1). The latter was used to define the borders of the BA. After several washing steps, the sections were mounted and coverslipped with Vectashield (Vector Laboratories). Multichannel confocal images of the LA and BA were obtained using a Nikon A1R microscope, apochromatic lens (CFI Plan Apo VC20X N.A. 1.40)(z stacks, 1 µm step size). Quantitative analyses were performed on a computer assisted image analysis system consisting of a MBF MS-88 computer-controlled motorized stage, a MBF DV-47d video camera, a Windows 7 PC computer, and StereoInvestigator program (MicroBrightField, Wiliston, VT), a custom designed morphology and stereology software. Tracings were made from the ventral through the dorsal extent of the amygdala. A total of six or seven tracings were made per amygdala, per hemisphere in each animal on both sides. After outlining the boundary of the LA and BA for each section at a low magnification, the software placed a set of optical dissector frames (50 x 50 µm) within each boundary in a systematic-random fashion where sampling grid was in the amygdala (grid size, 75 x 85 µm in the LA and 75 x 130 µm in the BA). Neurons were then counted in depth of 9 µm, according to stereological principles (West et al., 1991). In each amygdala, at least 500 neurons were sampled to ensure robustness of the data (Schmitz and Hof, 2000). Only NeuN+ cells were counted. We have noticed that some ZsGreen1-expressing cells with small soma (2-6% of all ZsGreen1-labeled cells) lacked NeuN staining, therefore were excluded from the counting. In addition, dense islands composed of 5-20 ZsGreen1-expressing small NeuN+ neurons were also excluded from the counting, as these neurons belong to the intercalated cell mass (see images taken at -1.85 and -2.1 mm to Bregma in Figure 1A).

**Figure 1.**
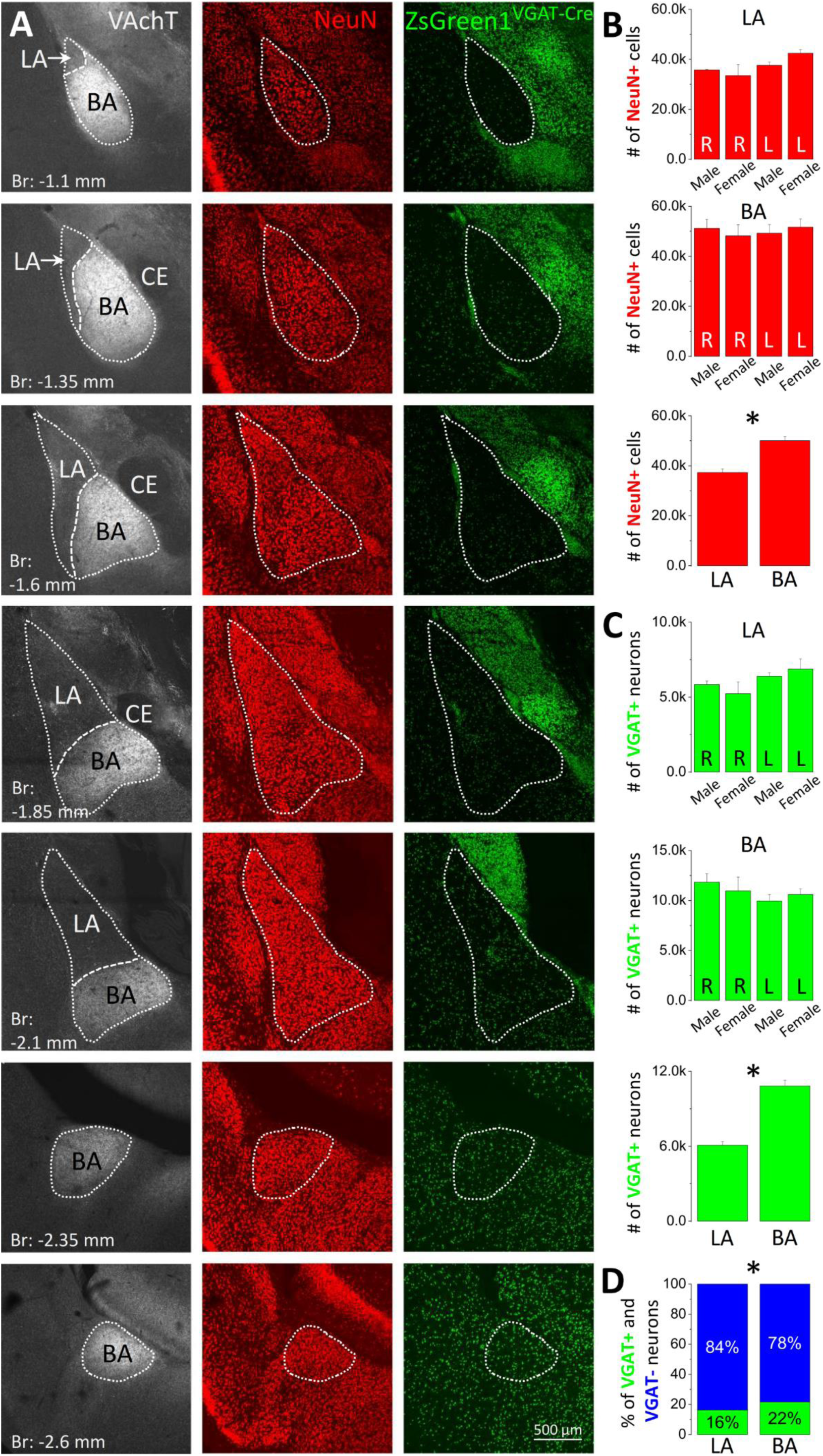
Total number of GABAergic and non-GABAergic neurons estimated in the lateral (LA) and basal nucleus (BA) of the mouse amygdala. ***A***, Serial sections containing the two amygdalar nuclei were prepared from an offspring of a Vgat-Cre mouse crossed with an Ai6 reporter mouse. In offspring, ZsGreen1, the reporter protein in Ai6 line is expressed in GABAergic neurons. NeuN+ cells lacking ZsGreen1 signal were consider non-GABAergic neurons. The borders of LA and BA were defined on the basis of NeuN immunostaining and vesicular acetylcholine transporter (VAchT), as VAchT-immunopositive axons are more abundant in the BA than in surrounding structures. Images were taken at the corresponding anterior-posterior coordinates relative to Bregma (Br). CE, central nucleus of the amygdala. ***B***, Using unbiased stereology, the total number of neurons expressing NeuN including both ZsGreen1-containing and lacking cells were assessed in the LA and BA of both sexes. No difference was observed between the hemispheres and/or sexes in the two nuclei (R, right; L, Left). In contrast, significantly more neurons were estimated in the BA in comparison to the LA (*, *p* < 0.001). ***C***, Similarly, no difference in the number of GABAergic (i.e. VGAT-expressing) neurons was observed between the hemispheres and/or sexes in the two nuclei. In contrast, the number of GABAergic cells was significantly larger in the BA (*, *p* < 0.001). ***D***, The ratio of GABAergic neurons between the two amygdalar nuclei was also significantly different (*, *p* < 0.001).

To determine the completeness and specificity of labeling in offspring generated by crossing Vgat-Cre with Ai6 mice, we immunostained sections with a cocktail of antibodies with an aim to visualize the all major inhibitory cell types (Table 1). Our counting revealed that ZsGreen1 was present in almost all GABAergic neurons identified with immunostaining (99.1 %, n = 1291, n = 2 mice), indicating that all known inhibitory cells are readily visualized in offspring. Then we counted how many ZsGreen1-expressing cells were also immunolabeled, which may be indicative for specificity. We found that the vast majority of ZsGreen1-containing NeuN+ neurons showed immunoreactivity for the mixture of GABAergic cell markers (86.5 %, n = 1480). The lack of immunostaining in 13.5 % of ZsGreen1-expressing neurons may be twofold. First, some GABAergic cells express neurochemical markers at a level below detectability using the applied method. That was the primary reason why we used distinct Cre mouse lines to visualize distinct groups of GABAergic cells in the subsequent experiments. Second, there may be a group of GABAergic cells (e.g. those expressing M2 type of muscarinic acetylcholine receptors) that can be only partially labeled by the cocktail of antibodies. These results collectively show that in Vgat-IRES-Cre mice, all major GABAergic cell types can be entirely labeled and non-GABAergic neurons are likely to be visualized only marginally, in line with earlier studies (Yamamoto et al., 2018).

### Surgical procedures and viral vectors

Anesthesia was induced and maintained with ketamine/xylazine anesthesia. Mice were secured in a stereotaxic frame and four injections per animal were aimed unilaterally at the following coordinates: 1.5 mm to Bregma (AP), 3.2 mm lateral to the midline (ML), 4.0 mm deep from the cortical surface (DV); 1.5 mm AP, 3.2 mm ML, 5.0 mm DV; 2.1 mm AP, 3.2 mm ML, 4.0 mm DV; and 2.1 mm AP, 3.2 mm ML, 5.0 mm DV. Adeno-associated virus (AAV)-based constructs engineered to transfect Cre+ and Cre+/Flp+ neurons with AAV2/5-EF1a-DIO-EYFP-WPRE-hGH and AAVdj-hSyn-C(on)/F(on)-EYFP-WPRE, respectively, were obtained from the University of Pennsylvania Vector Core and the University of North Carolina Vector Core, respectively. The virus titers were 3-6 x 10e12 vg/mL. At each site 350 nl (total of 1,400 nl/hemisphere) of AAV2/5-EF1a-DIO-EYFP-WPRE-hGH (flow rate: 50 nl/min) was unilaterally injected into the right BLA of 9-12 weeks old homozygous Pvalb-IRES-Cre, Sst-IRES-Cre and Vip-IRES-Cre mice. To visualize SST+ projection neurons in the amygdalar region, 300 nl of AAVretro-EF1a-mCherry-IRES-Flpo obtained from Addgene (titer, 7 x 10e12 vg/mL) was unilaterally injected in the basal forebrain (0.25 mm AP, 1.3 mm ML, 4.4 mm DV) or entorhinal cortex (4.25 mm AP, 3.25 mm ML, 3.5 mm DV) of Sst-IRES-Cre mice, followed by the injection of AAVdj-hSyn-C(on)/F(on)-EYFP-WPRE into the BLA at two AP coordinates as above (total of 400 nl/amygdala). In the case of three Npy^Cre^;Dlx5/6^Flp^ mice, AAVdj-hSyn-C(on)/F(on)-EYFP-WPRE (total of 1,400 nl/hemisphere) was injected into the amygdala using the same coordinates as above. In spite of the fact that the same amount of AAVdj was injected into Npy^Cre^;Dlx5/6^Flp^ mice, the spread of the viral infection was smaller as in case of AAV2/5 injection and only the LA was fully infected. Therefore, three additional Npy^Cre^;Dlx5/6^Flp^ mice were injected with ML coordinates modified from 3.2 mm to 2.8 mm, which resulted in full infection of the BA. The injection cannula was slowly withdrawn 5 minutes after injection. EYFP expression was allowed for four-five weeks, before the animals were sacrificed.

### Immunostaining of perfused tissue

Tissue samples for immunostaining from virus-injected and transgenic mice were prepared as for the stereological analysis. To estimate the ratios of inhibitory cell types, sections prepared from AAV-injected Cre mouse lines or transgenic mice were incubated in a mixture of primary antibodies, followed by a mixture of secondary antibodies listed in Table 1. To determine the proportion and the distribution of GABAergic boutons lacking immunoreactivity for PV, sections from virus-injected Pvalb-Cre mice were first pepsin-treated for antigen retrieval (Veres et al., 2014). Then, after several washing steps, the sections were incubated in guinea pig anti-VGAT, chicken anti-GFP, mouse anti-neuroligin 2, mouse anti-gephyrin and rabbit anti-Ankyrin G, followed by incubation in a mixture of DyL405-conjugated anti-guinea pig, A488-conjugated anti-chicken, Cy3-conjugated anti-mouse, A647-conjugated anti-rabbit. To reveal the neurotransmitter characteristics of axons in the contralateral amygdala upon injection of viral vectors into the amygdala region of NPY-Cre mice, immunostaining using a mixture of goat anti-GFP and rabbit anti-VGluT1 was perform. To visualize these antibodies, a mixture of Alexa488-conjugated anti-goat and Cy3-conjugated anti-rabbit was used. After several washing, the sections were mounted and coverslipped with Vectashield (Vector Laboratories) in each case. Multichannel confocal images were obtained using a Nikon A1R or C2 microscope, apochromatic lens (CFI Plan Apo VC20X N.A. 1.40)(z stacks, 1 µm step size). The image analysis was performed using Neurolucida Explorer.

### Electrophysiological slice recordings

Adult (P60-90) Npy-Cre x Ai14, Vgat^Cre^;CCK-GFPcoIN, Vip^Cre^;CCK-GFPcoIN or Sst-IRES-Cre and Npy^Cre^;Dlx5/6^Flp^ mice after four to six weeks following injection of viral vectors were deeply anesthetized with isoflurane, the brain was quickly removed and placed into ice-cold solution containing (in mM): 252 sucrose, 2.5 KCl, 26 NaHCO_3_, 0.5 CaCl_2_, 5 MgCl_2_, 1.25 NaH_2_PO_4_, 10 glucose, bubbled with 95% O_2_/5% CO_2_ (carbogen gas). Horizontal 200-µm thick brain sections containing the BLA were prepared with a vibratome (VT1200S, Leica Microsystems) and kept in an interface-type holding chamber containing ACSF at 36°C that gradually cooled down to room temperature. ACSF contained the followings (in mM): 126 NaCl, 2.5 KCl, 1.25 NaH_2_PO_4_, 2 MgCl_2_, 2 CaCl_2_, 26 NaHCO_3_, and 10 glucose, bubbled with carbogen gas. After at least a 60-minute long incubation, slices were transferred to a submerged-type recording chamber and perfused with 32-34°C ACSF with a flow rate of 1.5-2 mL/min.

Recordings were performed under visual guidance using differential interference contrast microscopy (via a model FN-1 Nikon or BX61W Olympus upright microscope) using a 40x water dipping objective. Fluorescent protein expression in neurons was visualized with the aid of a mercury arc lamp and a CCD camera (Andor Technology, Belfast, UK). Patch pipettes (5-7 MΩ) for whole-cell recordings were pulled from borosilicate capillaries with inner filament (thin walled, OD 1.5) using a P1000 pipette puller (Sutter Instrument, Novato, CA, USA). In whole-cell recordings the patch pipette contained a K-gluconate based intra-pipette solution (in mM): 115 K-gluconate, 4 NaCl, 2 Mg-ATP, 20 HEPES, 0.1 EGTA, 0.3 GTP (sodium salt), and 10 phosphocreatine adjusted to pH 7.3 using KOH, with an osmolarity of 290 mOsm/L. The pipette also contained 0.2% biocytin. Recordings were performed with a Multiclamp 700B amplifier (Molecular Devices, San Jose, CA, USA), low-pass filtered at 3 kHz, digitized at 10 kHz, recorded with an in-house data acquisition and stimulus software (Stimulog, courtesy of Prof. Zoltán Nusser, Institute of Experimental Medicine, Budapest, Hungary) or Clampex 10.4 (Molecular Devices), and were analyzed with EVAN 1.3 (courtesy of Professor Istvan Mody, Department of Neurology and Physiology, University of California, Los Angeles, CA), Clampfit 10.4 (Molecular Devices) and OriginPro 2018 (OriginLab Corp, Northampton, MA, USA).

For firing pattern analysis, neurons were recorded in current clamp mode at a holding potential of -65 mV. Voltage responses were tested with a series of hyperpolarizing and depolarizing square pulses of current with 800-ms duration and amplitudes between -100 and 100 pA at 10 pA step intervals, then up to 300 pA at 50 pA step intervals and finally up to 600 pA at 100 pA step intervals.

### Identification of recorded interneuron types and immunostaining in slices

Biocytin content of recorded neurons was visualized using Cy3-conjugated streptavidin in slices prepared from Vgat^Cre^;CCK-GFPcoIN Vip^Cre^;CCK-GFPcoIN and Npy^Cre^;Dlx5/6^Flp^ mice. Alexa488-conjugated and Alexa647-conjugated streptavidin was used to reveal the biocytin-loaded neurons in slices prepared from Npy-Cre x Ai14 mice and Sst-IRES-Cre mice, respectively. After the visualization of recorded neurons, confocal images of the filled cells were obtained using a confocal microscope (Nikon model C2) under a Plan-Apochromat VC 20x objective (N.A. 0.75, z step size: 1 µm, xy: 0.31 µm/pixel). Slices (not re-sectioned) were immunostained with antibodies based on the firing pattern characteristics and features of the dendritic and axonal arbors of the recorded neurons. Incubation of antibodies was performed for 7-8 days at 4 °C. Putative CCK+/CB1+ basket cells were immunostained with rabbit anti-CB1 antibody and visualized using DyL405-conjugated donkey anti-rabbit antibody. VIP content was tested by rabbit anti-VIP and visualized using Cy3-conjugated donkey anti-rabbit. To reveal the GFP content of CB1+ axon terminals in Vgat^Cre^;CCK-GFPcoIN mice, sections were incubated in a mixture of chicken anti-GFP and rabbit anti-CB1, followed by the visualization of these antibodies using Alexa488-conjugated donkey anti-chicken and Cy3-conjugated donkey anti-rabbit. The presence of nNOS and PV in SST+ inhibitory cells was revealed with goat anti-nNOS and rabbit anti-PV using DyL405-conjugated donkey anti-goat first and subsequently using donkey anti-rabbit in two cases, where no nNOS immunoreactivity was seen in the soma of tested neurons. To visualize the Kv2.1 type of voltage-gated potassium channels in slices, mouse anti-Kv2.1 antibody was used, which was developed by Cy3-conjugated donkey anti-mouse antibody.

The neurochemical content of biocytin-filled NPY+ interneurons was tested with the use of the following primary antibodies: rabbit anti-PV, guinea pig anti-SST, goat anti-nNOS, chicken anti-calbindin, guinea pig anti-calbindin, or rabbit anti-CB1. The following secondary antibodies were used to visualize these primary antibodies: DyL405-conjugated donkey anti-rabbit, DyL405-conjugated donkey anti-guinea pig, DyL405-conjugated donkey anti-chicken, DyL405-conjugated donkey anti-goat, Alexa647-conjugated donkey anti-rabbit, Cy5-conjugated donkey anti-goat or Cy5-conjugated donkey anti-guinea pig.

To reveal the axon initial segments, rabbit anti-Ankyrin G was used after antigen retrieval (Veres et al., 2014). This antibody was visualized with Alexa647-conjugated donkey anti-rabbit. All images were obtained using a confocal microscope (Nikon model C2) under a Plan Apo VC 60x objective (N.A. 1.4, z step size: 0.15-0.2 µm, xy: 0.08-0.10 µm/pixel). For the quantification of the input on axon initial segments, the images were subsequently de-convolved with Huygens software (SVI), and analyzed using the “Cell counter” and “SNT” plugins in the ImageJ software.

All antibodies used in this study are listed in Tables 2 and 3.

**Table 2.**
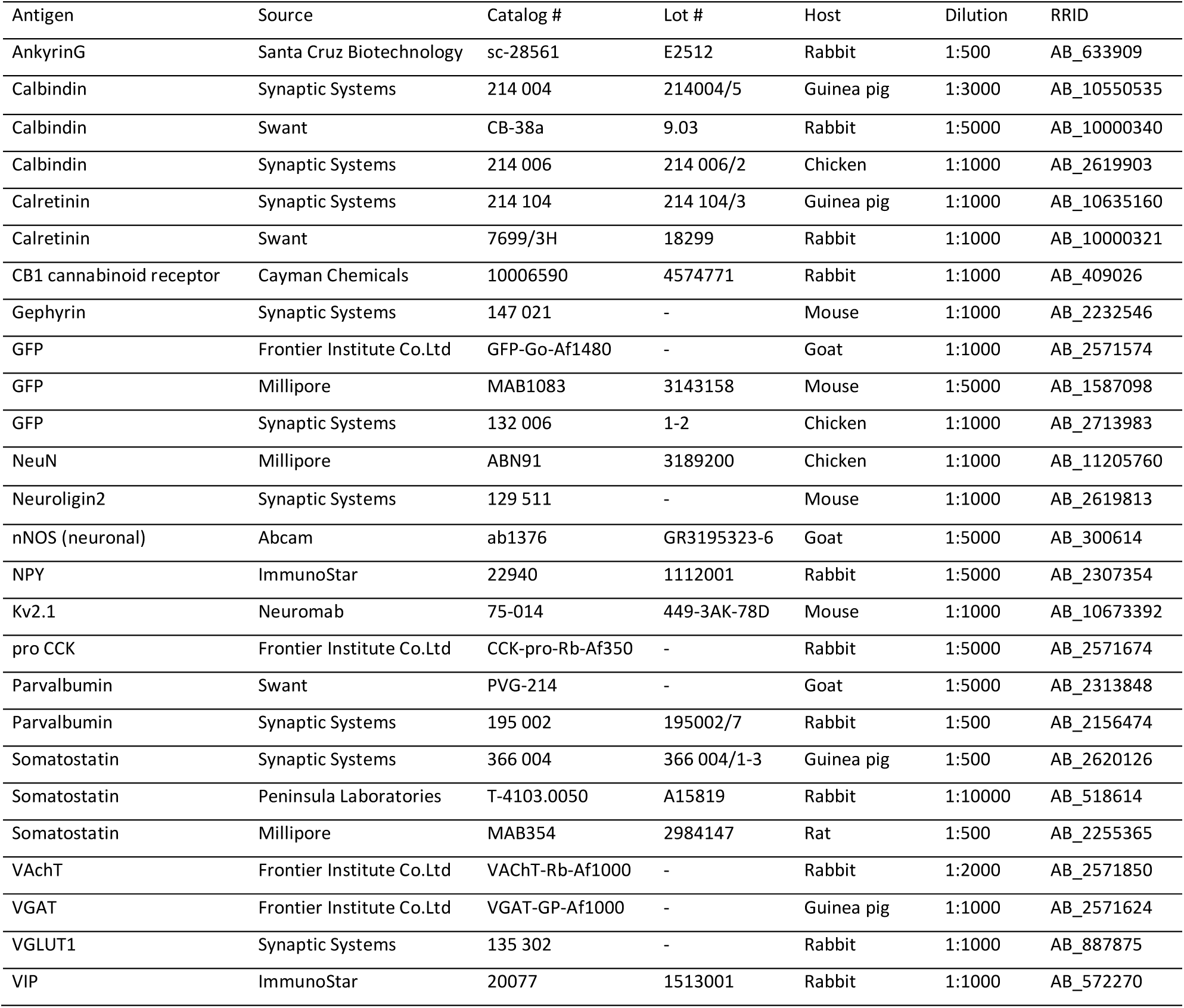
Primary antibodies used in this study

**Table 3.**
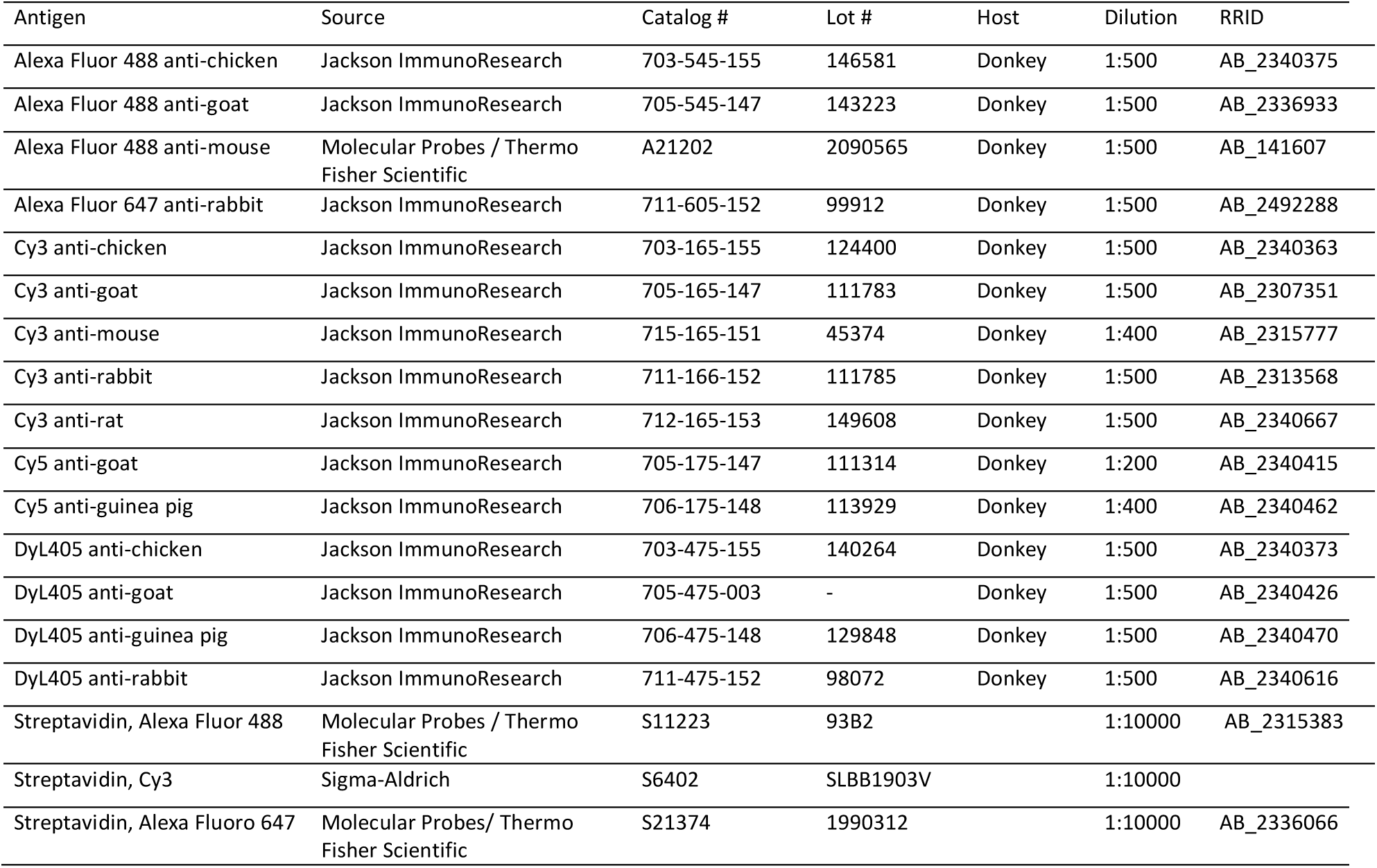
Secondary antibodies used in this study

| Antigen | Source | Catalog # | Lot # | Host | Dilution | RRID |
| --- | --- | --- | --- | --- | --- | --- |
| Alexa Fluor 488 anti-chicken | Jackson ImmunoResearch | 703-545-155 | 146581 | Donkey | 1:500 | AB_2340375 |
| Alexa Fluor 488 anti-goat | Jackson ImmunoResearch | 705-545-147 | 143223 | Donkey | 1:500 | AB_2336933 |
| Alexa Fluor 488 anti-mouse | Molecular Probes / Thermo Fisher Scientific | A21202 | 2090565 | Donkey | 1:500 | AB_141607 |
| Alexa Fluor 647 anti-rabbit | Jackson ImmunoResearch | 711-605-152 | 99912 | Donkey | 1:500 | AB_2492288 |
| Cy3 anti-chicken | Jackson ImmunoResearch | 703-165-155 | 124400 | Donkey | 1:500 | AB_2340363 |
| Cy3 anti-goat | Jackson ImmunoResearch | 705-165-147 | 111783 | Donkey | 1:500 | AB_2307351 |
| Cy3 anti-mouse | Jackson ImmunoResearch | 715-165-151 | 45374 | Donkey | 1:400 | AB_2315777 |
| Cy3 anti-rabbit | Jackson ImmunoResearch | 711-166-152 | 111785 | Donkey | 1:500 | AB_2313568 |
| Cy3 anti-rat | Jackson ImmunoResearch | 712-165-153 | 149608 | Donkey | 1:500 | AB_2340667 |
| Cy5 anti-goat | Jackson ImmunoResearch | 705-175-147 | 111314 | Donkey | 1:200 | AB_2340415 |
| Cy5 anti-guinea pig | Jackson ImmunoResearch | 706-175-148 | 113929 | Donkey | 1:400 | AB_2340462 |
| DyL405 anti-chicken | Jackson ImmunoResearch | 703-475-155 | 140264 | Donkey | 1:500 | AB_2340373 |
| DyL405 anti-goat | Jackson ImmunoResearch | 705-475-003 | - | Donkey | 1:500 | AB_2340426 |
| DyL405 anti-guinea pig | Jackson ImmunoResearch | 706-475-148 | 129848 | Donkey | 1:500 | AB_2340470 |
| DyL405 anti-rabbit | Jackson ImmunoResearch | 711-475-152 | 98072 | Donkey | 1:500 | AB_2340616 |
| Streptavidin, Alexa Fluor 488 | Molecular Probes / Thermo Fisher Scientific | S11223 | 93B2 |  | 1:10000 | AB_2315383 |
| Streptavidin, Cy3 | Sigma-Aldrich | S6402 | SLBB1903V |  | 1:10000 |  |
| Streptavidin, Alexa Fluoro 647 | Molecular Probes/ Thermo Fisher Scientific | S21374 | 1990312 |  | 1:10000 | AB_2336066 |

### Reconstruction of biocytin-loaded cells

The dendritic and axonal arbors of the intracellularly-filled neurons were reconstructed with Neurolucida 10.53 software, using confocal stacks acquired of the cell. The drawings of each neuron were analyzed with Neurolucida Explorer, and the values were corrected for shrinkage and flattening of the tissue (correction factor in the z axis: 1.7. No correction in the x and y axis). Branched Structure Analysis was used to study the dendritic length and number of nodes. Sholl analysis was used to estimate the complexity of the dendritic arbor by determining the number of processes crossing concentric spheres centered on the cell soma with 50 µm increments in their radius. Close apposition of a labeled bouton onto its target was defined as no apparent gap between the two profiles in 3D view.

### Statistical analysis

Data are presented as mean ± SEM, if not indicated otherwise. Statistical significance (*p* < 0.05) was assessed by t-test for comparison of data with a normal distribution, whereas Kruskal-Wallis (KW) ANOVA, Dunn’s test, Mann-Whitey (MW) U test and Kolmogorov-Smirnov (KS) test were used for datasets with a non-normal distribution.

## Results

Using unbiased stereology we assessed the number of GABAergic neurons in the LA and BA. Our investigations were performed both in the right and left hemispheres of male and female mice. To estimate the total number of GABAergic and non-GABAergic neurons, we counted the number of NeuN+ cells expressing ZsGreen1 (i.e. GABAergic neurons) or lacking this reporter protein (i.e. non-GABAergic neurons) in the LA and BA of mice generated by breeding of Vgat-Cre and Ai6 mice (Figure 1, Table 4). Comparison of the number of all neurons between hemispheres revealed no difference: a similar number of neurons was found in the right and left LA (*p* = 0.29 for males, *p* = 0.17 for females) as well as in the right and left BA (*p* = 0.72 for males, *p* = 0.57 for females)(Figure 1B). In addition, we found no sex difference in the number of all neurons, when the corresponding amygdalar nuclei were compared (*p* = 0.67 for the right LA between males and females; *p* = 0.08 for the left LA between males and females; *p* = 0.63 for the right BA between males and females; *p* = 0.64 for the left BA between males and females)(Figure 1B). However, a larger number of neurons were found in the BA than LA (37,304 ± 1,439 in the LA; 50,044 ± 1,649 in the BA; n =12, pooled data from both males and females as well as from right and left hemispheres, *p* < 0.001)(Figure 1B).

**Table 4.** Estimation of the number of GABAergic (VGAT+/NeuN+) and all neurons (NeuN+), including both GABAergic and non-GABAergic neurons in the LA and BA using unbiased stereology.

|  |  | **LA** |  | **BA** |  |
| --- | --- | --- | --- | --- | --- |
|  | Animal ID | VGAT+<br>/NeuN+ # | All<br>NeuN+ # | VGAT+<br>/NeuN+ # | All<br>NeuN+ # |
| Male_right | Mouse # 1 | 5596 | 35337 | 11051 | 46925 |
|  | Mouse # 2 | 5585 | 35646 | 10937 | 48101 |
|  | Mouse # 3 | 6317 | 36180 | 13507 | 58462 |
| Female_right | Mouse # 4 | 3683 | 26025 | 9004 | 43899 |
|  | Mouse # 5 | 5834 | 32994 | 10214 | 43567 |
|  | Mouse # 6 | 6170 | 41439 | 13653 | 57025 |
| Male_left | Mouse # 1 | 6778 | 34971 | 8755 | 43398 |
|  | Mouse # 2 | 6469 | 38774 | 10962 | 55323 |
|  | Mouse # 3 | 5896 | 39040 | 10156 | 48941 |
| Female_left | Mouse # 4 | 8031 | 44230 | 9569 | 46222 |
|  | Mouse # 5 | 5655 | 39434 | 10809 | 51056 |
|  | Mouse # 6 | 6908 | 43578 | 11449 | 57605 |

Similarly, comparison of the number of GABAergic neurons between hemispheres revealed no difference, i.e. a comparable number of VGAT+ neurons was observed in the right and left LA (*p* = 0.20 for males, *p* = 0.19 for females) as well as in the right and left BA (*p* = 0.16 for males, *p* = 0.83 for females)(Figure 1C). Moreover, we have not found a sex difference in the number of GABAergic neurons, when the corresponding amygdalar nuclei were compared (*p* = 0.52 for the right LA between males and females; *p* = 0.56 for the left LA between males and females; *p* = 0.62 for the right BA between males and females; *p* = 0.49 for the left BA between males and females)(Figure 1C). Yet, a significantly larger number of VGAT+ neurons were identified in the BA than LA (6,077 ± 298 in the LA; 10,839 ± 441 in the BA; n = 12, *p* < 0.001)(Figure 1C). The larger difference in the number of GABAergic neurons between the LA and BA in comparison to the difference in the number of total neurons between these two nuclei predicted that the proportion of VGAT+ neurons in the neuronal populations should be different. Indeed, the ratio of GABAergic neurons in the BA was significantly larger than in the LA (16.3 ± 0.5 % in the LA; 21.6 ± 0.4 % in the BA; n = 12, *p* < 0.001)(Figure 1D). These data collectively revealed that the LA contains a lower number of neurons than the BA, and the fraction of GABAergic neurons in the LA is smaller than in the BA. Importantly, we observed no difference in the number of neurons either between hemispheres or sexes.

Next, we looked at whether the ratios of GABAergic cells in two amygdala nuclei at different anterior-posterior distances from Bregma show any difference (Table 5). In the LA, no substantial change in the ratio of inhibitory cells could be noticed (KW-ANOVA, *p* > 0.07). In contrast, there was a significant difference in the ratio of GABAergic cells in the BA at distinct anterior-posterior levels (KW-ANOVA, *p* < 0.001). Specifically, the proportion of GABAergic cells at the most posterior part of the BA (at -2.6 mm from Bregma) was significantly lower than those found at the distance of -1.85 and -2.1 mm. In addition, the ratio of GABAergic cells was also lower at -1.35 mm in comparison to that observed at -2.1 mm (Table 5). Interestingly, the portion of inhibitory neurons ‘peaked’ at -2.1 mm for both the LA and BA. These data suggest that the inhibition at distinct anterior-posterior distances, at least in the BA may be different.

**Table 5.** The ratio of GABAergic cells in the lateral (LA) and basal (BA) amygdala nuclei at different anterior-posterior distances from Bregma. ‘n’ refers to the number of sections used for the calculation.

|  | Distance from Bregma (mm) |  |  |  |  |  |  |
| --- | --- | --- | --- | --- | --- | --- | --- |
|  | -1.1 | -1.35 | -1.6 | -1.85 | -2.1 | -2.35 | -2.6 |
|  | % of GABAergic cells at different distances from Bregma |  |  |  |  |  |  |
| LA |  | 12.9 ± 1.1 | 14.6 ± 1.1 | 15.3 ± 1.2 | 17.8 ± 0.8 | 16.3 ± 2.3 |  |
|  |  | n=12 | n=12 | n=12 | n=12 | n=8 |  |
| BA | 20.8 ± 3.9 | 19.5 ± 1.2 | 22.5 ± 1.8 | 25.1 ± 0.9 | 25.9 ± 0.6 | 20.6 ± 0.9 | 16.5 ± 1.3 |
|  | n=10 | n=12 | n=12 | n=12 | n=12 | n=12 | n=11 |
|  | Normalized ratio of GABAergic cells at different distances from Bregma (%) |  |  |  |  |  |  |
| LA |  | 84.3 ± 7.4 | 94.5 ± 7 | 99.5 ± 7.9 | 115.5 ± 5.1 | 105.5 ± 15 |  |
| BA | 96.7 ± 18.5 | 90.4 ± 5.5<sup>a</sup> | 104.4 ± 8.2 | 116.6 ± 4.6<sup>b</sup> | 120.0 ± 2.9<sup>a,c</sup> | 95.4 ± 4.6 | 76.6 ± 5.9<sup>b,c</sup> |

<sup>a</sup>, $p = 0.03$ ; <sup>b</sup>, $p = 0.0022$ ; <sup>c</sup>, $p < 0.001$ (Dunn's test)

As GABAergic neurons can be divided into several functional categories in cortical networks (Kepecs and Fishell, 2014), we next aimed to estimate the ratio of these major inhibitory neuron types within the circuits of the LA and BA using viral techniques combined with immunocytochemistry.

### PV+ interneurons targeting the perisomatic region of the principal cells

First, we assessed the proportion of PV-containing interneurons in the two amygdala nuclei by labeling PV+ interneurons in Pvalb-Cre mice using a viral vector (Figure 2A-C, Table 6). We found that significantly more PV+ interneurons were present in the BA than in the LA (Table 6). Previous studies uncovered that PV is expressed in two types of amygdalar interneurons that target the perisomatic region of principal neurons (Smith et al., 1998; Muller et al., 2006). PV+ basket cells expressing calbindin (CB) innervate the soma and the proximal dendrites of principal neurons, whereas PV+ axo-axonic cells lacking CB form synaptic contacts on axon initial segments (Bienvenu et al., 2012; Veres et al., 2014; Vereczki et al., 2016; Veres et al., 2017). Therefore, by using immunostaining against CB we separated these two PV-containing interneuron types and estimated their ratio (Figure 2D). In line with earlier findings (McDonald and Betette, 2001; McDonald and Mascagni, 2001), CB was present in the majority of PV+ interneurons (Figure 2E), i.e. PV+ basket cells are the most abundant interneurons among PV+ GABAergic neurons (Table 6). Based on our counting, PV+/CB+ basket cells constitute ∼ 2.2 % and ∼ 4.7 % of all neurons in the LA and BA, respectively.

**Figure 2.**
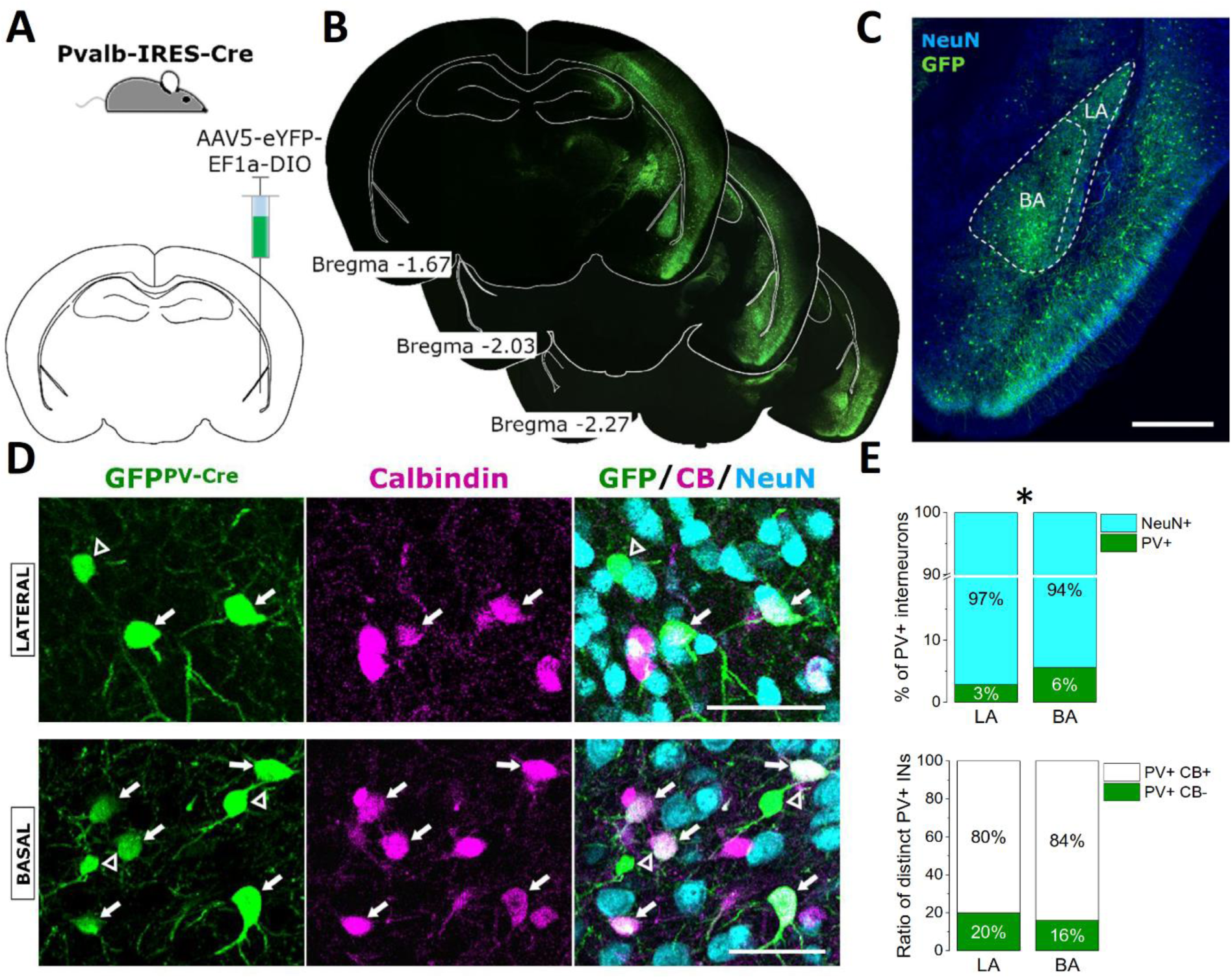
Ratio of parvalbumin (PV)-expressing interneurons in the LA and BA. ***A***, Schematic of the strategy for targeting PV-expressing interneurons in the amygdala. ***B***, Representative images of EYFP expression after virus transfection taken at the corresponding anterior-posterior coordinates (in mm) relative to Bregma. As EYFP labeling was enhanced by immunostaining with an antibody developed against GFP, we refer to the enhanced EYFP signal as GFP here and in the latter figures. ***C***, Representative example of the amygdalar region taken at a higher magnification (scale bar = 500 µm). ***D***, Majority of GFP-labeled neurons in Pvalb-Cre mice contains calbindin (CB)(arrows), whereas a minority of GFP-expressing interneurons lacks immunopositivity for this Ca^2+^ binding protein (open arrowheads)(scale bars = 50 µm). ***E***, The percentage of PV+ GABAergic cells in the LA and BA is different (top, *, *p* < 0.001). In contrast, there is no difference in the ratio of PV+ interneurons expressing CB in the two amygdalar nuclei (bottom).

**Table 6.** Fraction of distinct types of GABAergic cells in the LA and BA. In each case, the counting was performed in sections prepared from three mice. # of NeuN+ cells refers to the total number of neurons counted in the given number of sections (# of sec.) indicated. Significant differences are shown in bold.

| Neuron types | LA<br>% of inh.<br>neurons | # of<br>NeuN+<br>cells | # of<br>sec. | BA<br>% of inh.<br>neurons | # of<br>NeuN+<br>cells | # of<br>sec. | t test<br>*p* value |
| --- | --- | --- | --- | --- | --- | --- | --- |
| All PV+ | 2.8 ± 0.4 | 4,824 | 7 | 5.6 ± 0.3 | 8,101 | 12 | **< 0.001** |
| PV+Calb+ basket<br>cells | 80.1 ± 4.2% of<br>all PV+<br>interneurons |  |  | 84.1 ± 1.4% of<br>all PV+<br>interneurons |  |  | 0.39 |
| CCK+ GABAergic<br>cells in<br>Vgat<sup>Cre</sup>;CCK-<br>GFPcoIN | 1.1 ± 0.3 | 7,768 | 10 | 2.0 ± 0.2 | 8,815 | 12 | **0.0057** |
| proCCK+ in<br>Vgat<sup>Cre</sup>;CCK-<br>GFPcoIN | 53.4 ± 2.3 %<br>of all GFP+<br>interneurons |  |  | 62.8 ± 1.9 %<br>of all GFP+<br>interneurons |  |  | **0.024** |
| Interneuron-selective<br>interneurons | 4.8 ± 0.4 | 6,824 | 7 | 6.9 ± 0.26 | 8,772 | 12 | **0.0013** |
| All SST+<br>interneurons | 2.3 ± 0.3 | 6,031 | 10 | 5.4 ± 0.8 | 10,406 | 13 | **0.0018** |
| SST+nNOS+ | 41.6 ± 5.4% of<br>all SST+<br>neurons |  |  | 25 ± 5.4% of<br>all SST+<br>neurons |  |  | **0.043** |
| All NPY+<br>GABAergic cells | 3.8 ± 0.3 | 10,458 | 12 | 8.1 ± 0.7 | 6,931 | 11 | **< 0.001** |
| NPY+PV+ | 23.7 ± 1.9% of<br>all NPY+<br>GABAergic<br>cells |  |  | 29.7 ± 2.7% of<br>all NPY+<br>GABAergic<br>cells |  |  | 0.085 |
| NPY+SST+ | 30.9 ± 3.6% of<br>all NPY+<br>GABAergic<br>cells |  |  | 25.5 ± 3.0% of<br>all NPY+<br>GABAergic<br>cells |  |  | 0.26 |
| Neurogliaform cells | 46.6 ± 4.7 %<br>of all NPY+<br>GABAergic<br>cells |  |  | 42.9 ± 2.8 %<br>of all NPY+<br>GABAergic<br>cells |  |  | 0.51 |

### One third of axo-axonic cells lack PV

Previous studies have shown that some axo-axonic cells may lack PV in cortical areas (He et al., 2016; Paul et al., 2017), thus the ratio of axo-axonic cells determined solely on the basis of PV expression (LA: 19.9 ± 4.2 %; BA: 15.9 ± 1.4 % of all PV+ interneurons) could be underestimated in the amygdala as well. To test the presence of amygdalar axo-axonic cells lacking PV, we examined the PV content of GABAergic boutons forming synaptic contacts with axon initial segments. The presence of a synapse between a closely apposed bouton and an axon initial segment was assessed by visualizing two inhibitory synapse-specific proteins, gephyrin and neuroligin 2 using immunostaining (Veres et al., 2014). In virus-injected Pvalb-Cre mice, the GFP content of boutons was indicative for PV expression, whereas GABAergic phenotype of axon terminals was revealed by immunostaining against VGAT (Figure 3A). We observed that the minority (∼ 32 %) of VGAT+ axon terminals forming synaptic contacts with Ankyrin G-labeled axon initial segments originated from PV- axo-axonic cells in both amygdalar nuclei (LA: 32.2 ± 2.8 %, BA: 32.1 ± 1.7 %, n=14-14 axon initial segments, 2 animals)(Figure 3B). If we assume that PV+ and PV- axo-axonic cells in the LA and BA give rise to a similar number of axonal boutons and contact the individual axon initial segments with a similar number of synapses, then approximately 30 % of this inhibitory cell type cannot be labeled in Pvalb-Cre mice. By summing up the ratio of PV+/CB- interneurons (i.e. PV+ axo-axonic cells) and PV- axo-axonic cells, the ratio of all axo-axonic cells can be 0.8 % of all neurons in the LA and 1.3 % in the BA.

**Figure 3.**
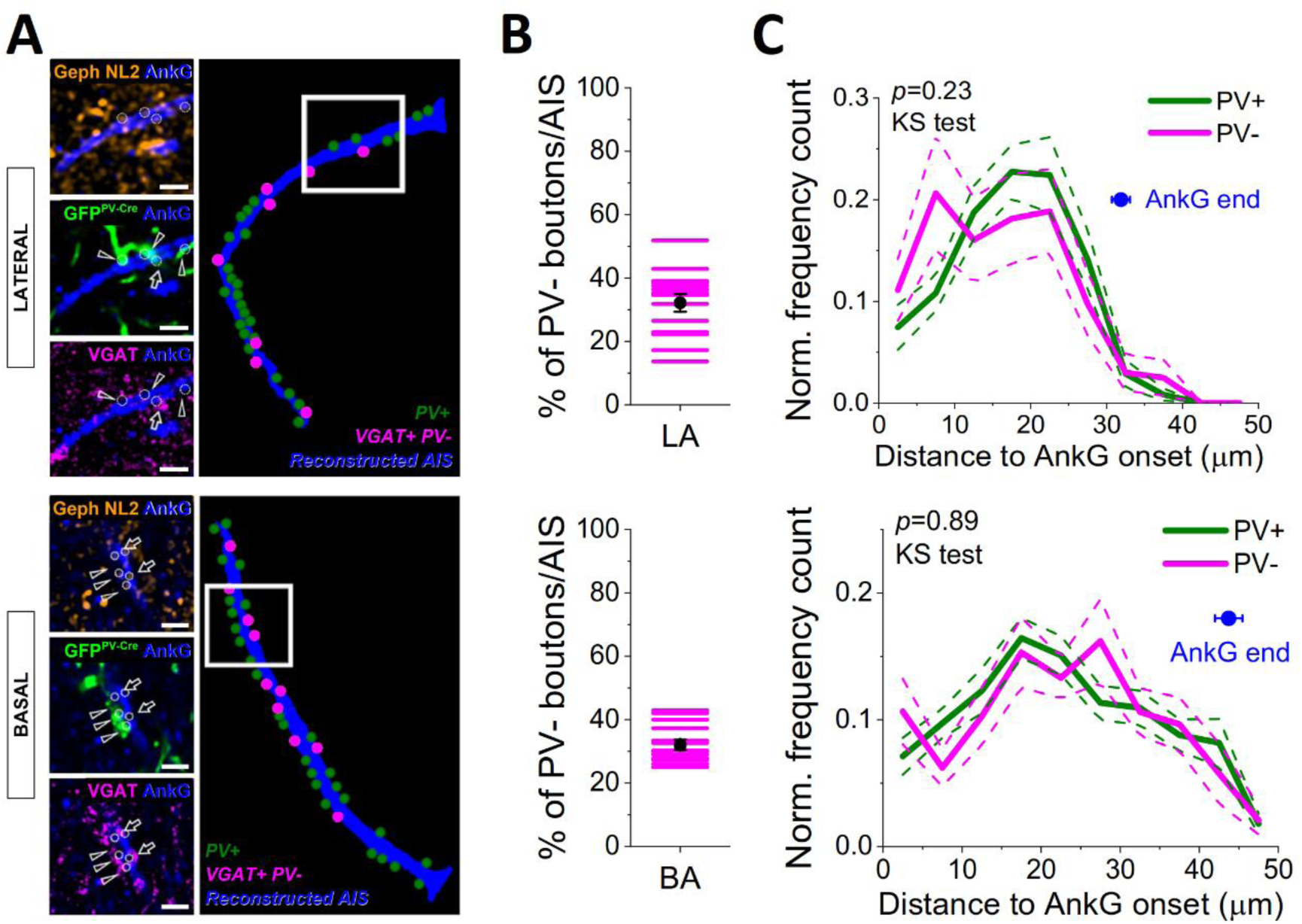
One third of GABAergic boutons apposing axon initial segments lack PV both in the LA and BA. ***A***, Examples of axon initial segments receiving synaptic contacts from both PV+ (arrowhead) and PV- boutons (arrows). PV-expressing boutons were visualized with GFP in Pvalb-Cre mice using a viral vector, whereas PV-immunonegative axon terminals were determined by immunostaining against VGAT. Synaptic contacts between the boutons and Ankyrin G (AnkG)-labeled axon initial segments were revealed by the presence of two inhibitory synapse-specific proteins, gephyrin (Geph) and neuroligin 2 (NL2)(scale bars = 2 µm). ***B***, A substantial portion of GABAergic boutons forming synaptic contacts with axon initial segments lacked GFP content, indicative for the absence of PV expression (n = 14-14 axon initial segments in the LA and BA were reconstructed). ***C***, The spatial distribution of PV+ and PV- axon terminals along Ankyrin G-immunostained profiles was similar (KS test, Kolmogorov-Smirnov test), indicating that axo-axonic cells irrespective of their PV content prefer to innervate a given portion of axon initial segments. Mean, solid line; S.E.M., dashed line. Blue dots indicate the average length of the reconstructed Ankyrin G-immunoreactive profiles.

Next, we asked whether PV- axo-axonic cells follow the same innervation strategy as PV+ axo-axonic cells. Our previous investigations revealed that PV+ axo-axonic cells in the BA prefer to innervate that part of the axon initial segments where the action potential generation has the highest likelihood (Veres et al., 2014). Therefore, we compared the number of PV+ and PV-/VGAT+ axon terminals along axon initial segments both in the LA and BA. We found that in both nuclei the spatial distribution of boutons originating from the two neurochemically different axo-axonic cells were similar (119 PV-/VGAT+ boutons and 251 PV+ boutons examined along 14 axon initial segments in the LA, *p* = 0.23; 196 PV-/VGAT+ boutons and 415 PV+ boutons examined along 14 axon initial segments in the BA, *p* = 0.89, Kolmogorov-Smirnov test, 2 mice)(Figure 3C). These results indicate that axo-axonic cells, irrespective of their PV content, preferentially target a given portion of axon initial segments both in the LA and BA. However, the preferentially targeted portion measured from the onset of the Ankyrin G-labeled profiles was different in the LA and BA (*p* < 0.0001, Kolmogorov-Smirnov test, Figure 3C). Of all GABAergic boutons along the axon initial segments (PV+ and PV- together), 60 % was found between 10-25 µm in the LA and 12-35 µm in the BA. Taking into account that the length of the Ankyrin G-immunostained profiles in the LA was significantly shorter than in the BA (LA: 31.9 ± 1.2 µm, n =14; BA: 43.7 ± 1.7 µm, n = 14; *p* < 0.001, Figure 3C), the relative onset and extent of that portion of axon initial segments, which was covered by the majority of inhibitory inputs, was rather similar in both nuclei. Specifically, 60% of all GABAergic boutons were spread between 31.3 % and 78.6 % of the length of axon initial segment in the LA and between 27.4 % and 80 % in the BA. These observations thus suggest that the relative length for spike generation along the axon initial segment densely covered by GABAergic boutons is rather similar in the LA and BA principal cells (Veres et al., 2014).

### CCK+/CB1+ basket cells

In addition to the axo-axonic cells and PV+ basket cells, basket cells expressing CCK and CB1 contribute substantially to GABAergic innervation of the perisomatic region of principal cells in the BA (Vereczki et al., 2016). To estimate the proportion of CCK+/CB1+ basket cells in the two amygdala nuclei, we generated a novel mouse line, BAC-CCK-GFPcoIN_sb in which the GFP expression in CCK+ neurons is controlled in a Cre-dependent manner. By crossing this new mouse line with Vgat-Cre mice, GFP expression in the LA and BA (and in other cortical regions) was found to be restricted to the CCK+ GABAergic cells in offspring, i.e. in Vgat^Cre^;CCK-GFPcoIN mice (Figure 4A). To uncover which interneuron types express GFP under CCK promoter in these mice, we performed whole-cell recordings in green neurons in slice preparations followed by neurochemical analysis of the recorded neurons (n = 38, 10 in the LA and 28 in the BA). Only those interneurons were included in the analysis that could be tested for immunoreactivity of CB1 and/or VIP. Our investigations revealed that the vast majority of GFP-expressing interneurons was CCK+/CB1+ basket cell (71.1 %, n = 38 recorded green neurons) as these interneurons had axon terminals immunoreactive for CB1 (27 out of 27)(Figure 4B). The minority of recorded neurons (28.9 %, n = 38 recorded green cells) was found to be immunonegative for CB1 (8 out of 8 tested for axon terminals), but showed immunopositivity for VIP in the soma (4 out of 4 tested)(Figure 4C). This latter interneuron population had small somata and resembled interneuron-selective interneurons based on their morphological appearance and spiking features (Figure 4D, Table 7, (Rhomberg et al., 2018)). To confirm that a portion of VIP+ interneuron-selective interneurons can be labeled in the BAC-CCK-GFPcoIN_sb mouse line, we crossed these mice with Vip-Cre mice. In their offspring, i.e. in Vip^Cre^;CCK-GFPcoIN mice, we observed GFP+ interneurons with small somata that were immunopositive for VIP (LA: 95 %, n = 64; BA: 96 %, n = 75, 3 mice), but lacked immunoreactivity for proCCK (LA: 96 %, n = 50; BA: 98.7 %, n = 78, 3 mice)(Figure 4E, F). We performed whole-cell recordings from small GFP+ interneurons in slices prepared from in Vip^Cre^;CCK-GFPcoIN mice. Both the morphological appearance, including short dendrites and locally ramified axon collaterals and single-cell features of recorded interneurons (n = 5) were similar to those small green interneurons that were sampled in Vgat^Cre^;CCK-GFPcoIN mice (Figure 4G, Table 7). Importantly, both the morphological appearance and firing properties of small green interneurons recorded either in Vgat^Cre^;CCK-GFPcoIN mice or Vip^Cre^;CCK-GFPcoIN mice were comparable to those reported earlier for VIP+ interneurons in the amygdala (Rhomberg et al., 2018). In summary, these results collectively suggest that in Vgat^Cre^;CCK-GFPcoIN mice, CCK+/CB1+ basket cells with large somata are labeled predominantly, while GFP+ interneurons with small somata are VIP+ interneuron-selective interneurons. Thus, by using proCCK antibody, which does not stain VIP+ interneuron-selective interneurons under our circumstances, the number of CCK+/CB1+ basket cells can be estimated accurately in Vgat^Cre^;CCK-GFPcoIN mice.

**Figure 4.**
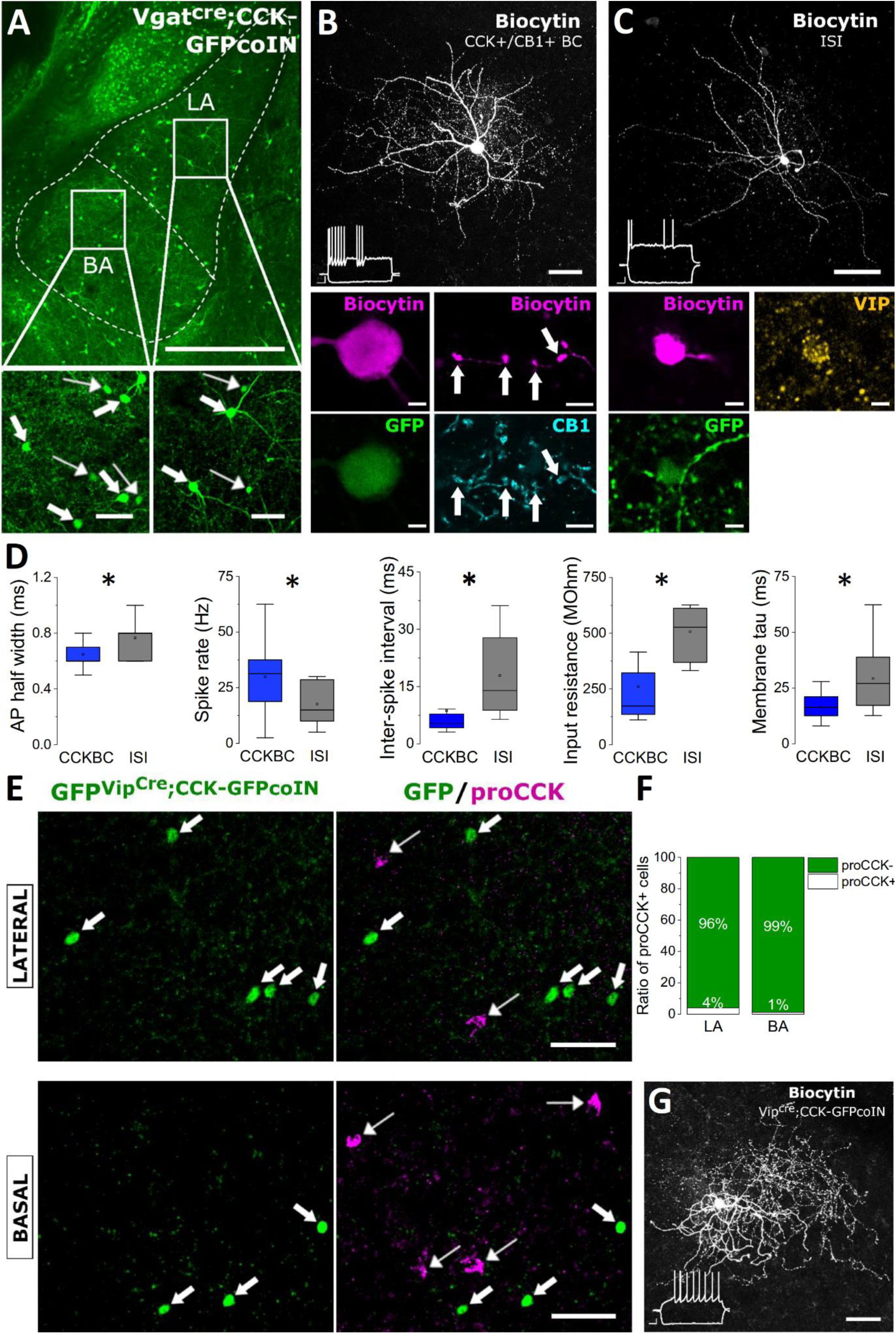
Characterization of interneurons expressing GFP in the amygdala of Vgat^Cre^;CCK-GFPcoIN and Vip^Cre^;CCK-GFPcoIN mice. ***A***, In Vgat^Cre^;CCK-GFPcoIN mice, GFP expression visualizes GABAergic cells with large (thick arrows) and small (thin arrows) somata in both amygdalar nuclei (scale bars = 500 µm and 50 µm for large and small images, respectively). ***B***, An example GFP-expressing interneuron with large soma recorded in an acute slice prepared from a Vgat^Cre^;CCK-GFPcoIN mouse (scale bar = 50 µm). The interneuron was identified as CCK+/CB1+ basket cell (BC) based on its firing pattern and the CB1 immunoreactivity in its axonal boutons (white arrows in small images)(scale bar = 5 µm). ***C***, An example for a GFP-expressing interneuron with small soma (scale bar = 50 µm). The interneuron was identified as interneuron-selective interneuron (ISI) based on its firing characteristics and VIP content in its soma (scale bar = 5 µm). ***D***, Differences in single-cell features between CCK+/CB1+ basket cells (CCKBC) and interneuron-selective interneurons (ISI) obtained *in vitro*. * labels significant difference (*p* < 0.05). Here and in Figure 8E, the mean (small open square), median (midline of the box), the interquartile range (box) and the 5/95 % values (whiskers) are shown on the charts. For data, see Table 7. ***E***, In the LA and BA of Vip^Cre^;CCK-GFPcoIN mice, GFP is expressed in interneurons with small somata (thick arrows) that are immunonegative for pro-cholecystokinin (proCCK, think arrows) (***F***) (scale bar = 50 µm). ***G***, An example interneuron recorded and labeled in an amygdalar slice (scale bar = 50 µm). Its voltage responses upon current injections (inset) are typical for VIP+ interneuron-selective interneurons. Scale bars of the firing patterns are x = 100 ms, y = 10 mV.

**Table 7.** Single-cell properties of CCK-expressing GABAergic cell types in the LA and BA. Data are presented as the median with the first and third quartiles in parentheses. P values obtained by statistical tests are indicated. Significant differences shown in bold and italic were determined by Kruskal-Wallis ANOVA and Mann-Whitney (MW) U test, respectively. CCKBC (CCK and CB1-expressing basket cell) and ISI (interneuron-selective interneurons) were recorded in Vgat^Cre^;CCK-GFPcoIN mice. VIP+/CCK+ ISI (interneuron-selective interneurons) were recorded in Vip^Cre^;CCK-GFPcoIN mice.

| Parameters | CCKBC<br>(n=25-27) | ISI<br>(n=7-8) | VIP+/CCK+ ISI<br>(n=5) | Kruskal-<br>Wallis<br>ANOVA | CCK<br>BC vs<br>ISI<br>MW<br>test | ISI vs<br>VIP+/<br>CCK+<br>MW<br>test |
| --- | --- | --- | --- | --- | --- | --- |
| AP half-width<br>(ms) | 0.6<br>(0.6, 0.7) | 0.8<br>(0.6, 0.8) | 0.7<br>(0.7, 0.9) | **0.036** | *0.037* | 0.84 |
| Spike rate (Hz) | 31.3<br>(18.8, 37.5) | 15<br>(10, 28.6) | 13.8<br>(11.3, 22.5) | **0.049** | *0.035* | 1 |
| AHP 50% decay<br>(ms) | 10.2<br>(5.4, 14.4) | 19.75<br>(8.45, 29.75) | 44.4<br>(22.6, 54.3) | **0.037\*** | 0.14 | 0.12 |
| ISI between the<br>first two spikes<br>(ms) | 5.3<br>(4.2, 7.8) | 13.95<br>(8.8, 27.8) | 22.1<br>(12.8, 25.7) | **0.002** | *0.005* | 0.93 |
| ISI between the<br>last two spikes<br>(ms) | 26.85<br>(22.6, 32.8) | 34.85<br>(5, 65.9) | 39.5<br>(33.6, 46.5) | 0.30 |  |  |
| Accommodation<br>ratio | 4.89<br>(3.92, 6.58) | 7.95<br>(2.76, 14.8) | 3.09<br>(1.81, 5.6) | 0.28 |  |  |
| Input resistance<br>(MΩ) | 173.45<br>(136.25, 321.8) | 527.2<br>(369.4, 612.1) | 359.6<br>(295.3, 457.8) | **0.005** | *0.002* | 0.19 |
| Membrane time<br>constant (ms) | 16.3<br>(12.55, 21.1) | 26.99<br>(17.19, 38.89) | 21.76<br>(19.87, 22.26) | **0.032** | *0.02* | 0.50 |
| Membrane<br>capacitance (pF) | 74.32<br>(51.19, 84.8) | 46.52<br>(39.05, 56.71) | 48.62<br>(36.73, 133.405) | 0.097 |  |  |
| Relative sag<br>amplitude | 0.14<br>(0.06, 0.24) | 0.09<br>(0.06, 0.16) | 0.11<br>(0.07, 0.14) | 0.402 |  |  |

AP, action potential; AHP, after hyperpolarization, ISI, inter-spike interval.
\*, there was a significant difference between CCKBC vs VIP+/CCK+ interneurons.

As a next step, we first assessed the ratio of CCK+ interneurons in the LA and BA in Vgat^Cre^;CCK-GFPcoIN mice (Figure 5A-C) and found significantly different number of these GABAergic cells in the two nuclei (Figure 5E, Table 6). Then, we estimated the fraction of CCK+/CB1+ basket cells within the population of CCK+ interneurons. As proCCK immunostaining readily labels basket cells, but not VIP+ interneuron-selective interneurons, we immunostained those sections that were used for calculation of the ratio of GFP+ interneurons in the amygdalar nuclei and counted the number of proCCK+ neurons (Figure 5D). However, we observed that there was a non-negligible amount of proCCK+ interneurons that were not labeled in the transgenic mice – 17 proCCK+/GFP- interneurons vs 116 GFP+ interneurons in the LA (10 sections, 3 mice) and 46 proCCK+/GFP- interneurons vs 180 GFP+ interneurons in the BA (12 sections, 3 mice)(Figure 5D). The perimeter of these proCCK+/GFP- interneurons was 45.4 ± 7.6 µm (n = 63), similar to the perimeter of the biocytin-labeled CCK+/CB1+ basket cells (48.1 ± 2.4 µm, n = 16, *p* = 0.3). Therefore, we assume that proCCK antibody used in our study labels CCK+/CB1+ basket cells in the amygdala, the majority of which expresses GFP (LA: 78.5 %, n = 79; BA: 71.1 %, n = 159).

**Figure 5.**
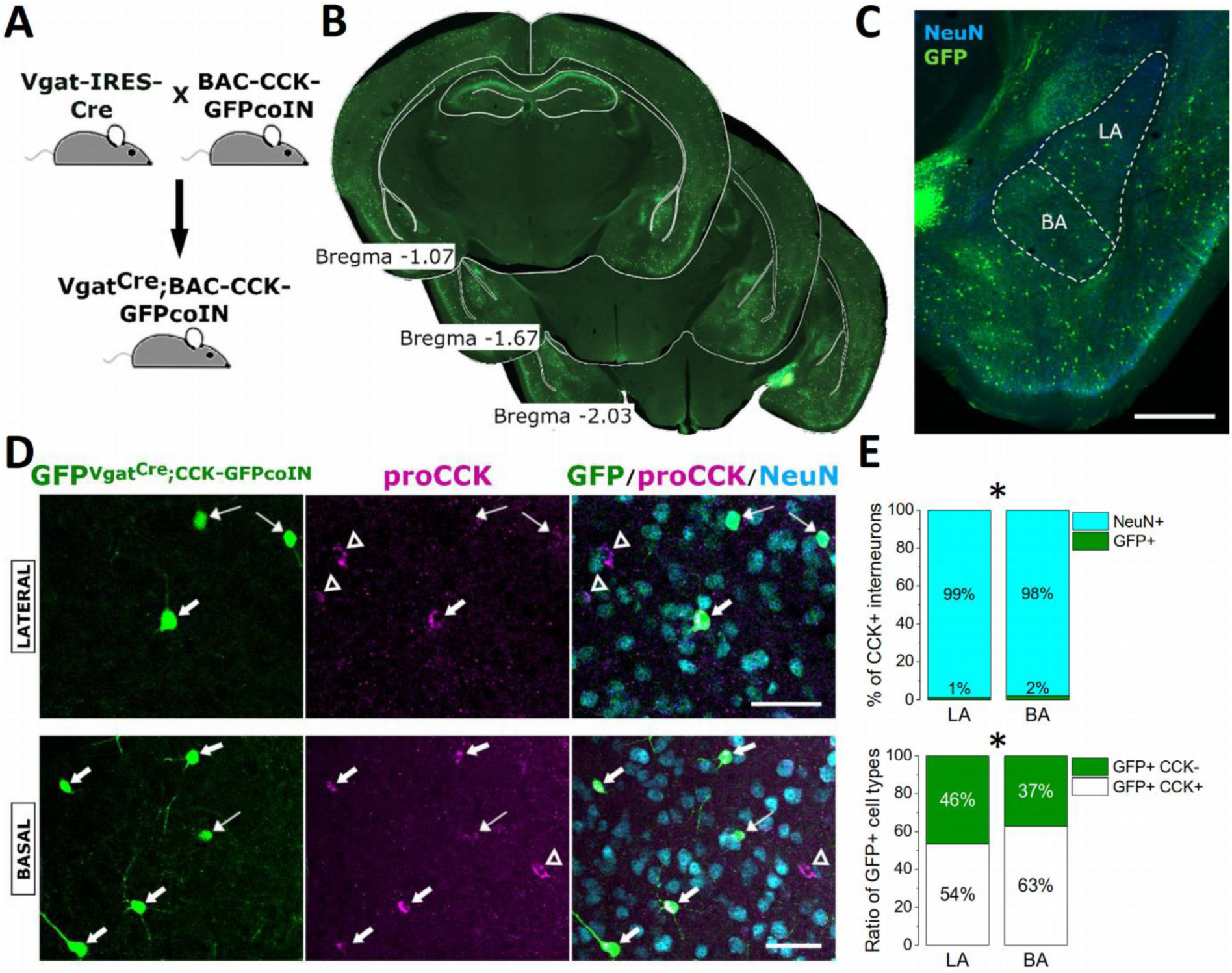
Number estimation for CCK+/CB1+ basket cells using proCCK immunostaining in Vgat^Cre^;CCK-GFPcoIN mice. ***A***, Strategy to visualize GABAergic neurons expressing CCK. ***B***, Representative images of GFP expression taken in a Vgat^Cre^;CCK-GFPcoIN mouse at the corresponding anterior-posterior coordinates (in mm) relative to Bregma. ***C***, Representative example of the amygdalar region taken at a higher magnification (scale bar = 500 µm). ***D***, A large portion of GFP-labeled neurons that have large somata in Vgat^Cre^;CCK-GFPcoIN mice is immunoreactive for pro-cholecystokinin (proCCK)(thick arrows), whereas a minority of GFP-expressing interneurons, typically with small somata, lacks immunopositivity for proCCK (thin arrows). A substantial number of proCCK+ GABAergic cells usually with large somata did not express GFP (open arrowheads)(scale bars = 50 µm). ***E,*** The percentage of CCK+ GABAergic cells in the LA and BA is different (top, *, *p* = 0.0057). In addition, there was a difference in the ratio of GFP+ interneurons expressing proCCK in the two amygdalar nuclei (bottom, *, *p* = 0.024).

To support the estimation of GFP expression in CCK+/CB1+ basket cells with an independent investigation, we evaluated the GFP content of the axon terminals originating from CCK+/CB1+ basket cells using immunostaining against CB1 in Vgat^Cre^;CCK-GFPcoIN mice. The analysis revealed that GFP signal was present in 63.5 ± 6.3 % and 66.7 ± 4.9 % of CB1+ boutons in the LA (n = 1,157 boutons, 13 sections, 4 mice) and in the BA (n = 2,025 boutons, 10 sections, 4 mice), respectively.

Based on these two distinct immunostainings, our results strongly suggest that ∼ 30-35 % of CCK+/CB1+ basket cells does not express GFP in Vgat^Cre^;CCK-GFPcoIN mice. Taking into account that a large portion, but not all of GFP+ interneurons are basket cells (LA: 53.4 %. BA: 62.8 %, Figure 5E, Table 6) and at least 1/3 of CCK+/CB1+ basket cells are not labeled in these transgenic mice, we propose that the ratio of this basket cell type is approximately 0.9 % in the LA and 1.9 % in the BA among all neurons.

### Interneuron-selective interneurons expressing VIP and/or CR

Next, we investigated the fraction of interneuron-selective interneurons in the LA and BA. Previous studies have established that VIP and CR are reliable markers for these GABAergic interneurons in cortical structures including the amygdala (Rhomberg et al., 2018; Krabbe et al., 2019). Therefore, we labeled VIP-expressing interneurons in Vip-Cre mice using a viral vector and visualized the CR content in neurons by immunostaining (Figure 6A-D). Our results show that interneuron-selective interneurons, including VIP+/CR+, VIP+/CR- and VIP-/CR+ subtypes, gave rise to a large portion of inhibitory cells (Figure 6E, Table 6). The ratio of interneuron-selective interneurons was significantly different between the two amygdalar nuclei (Table 6). In the LA, 35.9 ± 5.1 % of interneuron-selective interneurons expressed both VIP and CR, 31.9 ± 4.5 % expressed only VIP and 32.1 ± 4.7 % contained CR only. In the BA, 28.7 ± 3.2 % showed both VIP and CR labeling, 36.2 ± 3.5 % was labeled only for VIP and 35.1 ± 2.8 % was immunolabeled only for CR (Figure 6E). The fraction of the three interneuron-selective interneuron subtypes was comparable in both amygdalar nuclei (*p* > 0.22). These results show that VIP+ and/or CR+ interneuron-selective interneurons form the largest portion of GABAergic cell population in the LA and BA (Figure 11).

**Figure 6.**
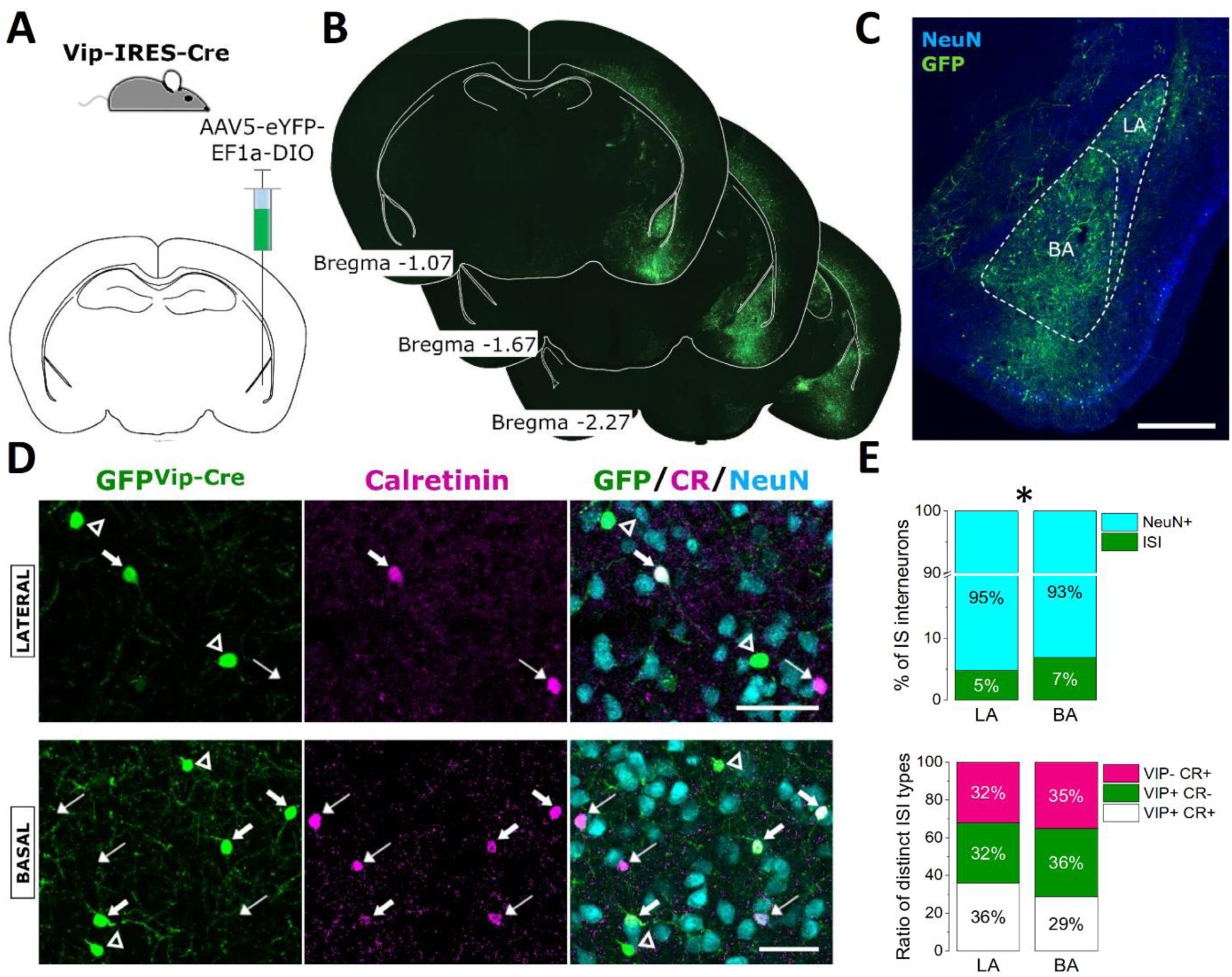
Interneuron-selective interneurons (ISI) in the LA and BA. ***A***, Schematic of the strategy for targeting VIP-expressing interneurons in the amygdala. ***B***, Representative images of GFP expression after virus transfection taken at the corresponding anterior-posterior coordinates (in mm) relative to Bregma. ***C***, Representative example of the amygdalar region taken at a higher magnification (scale bar = 500 µm). ***D***, A portion of GFP-labeled neurons in Vip-Cre mice contains calretinin (CR)(thick arrows), or lacks immunopositivity for this Ca^2+^ binding protein (open arrowheads), but there are also CR-immunoreactive somata that do not contain GFP (thin arrows). Note that all labeled interneurons had small somata irrespective of their neurochemical content (scale bars = 50 µm). ***E***, The percentage of interneuron-selective interneurons in the LA and BA differed (top, *, *p* = 0.0013). On the other hand, the ratios of the three neurochemically distinct ISI types (VIP+CR+, VIP+CR- and VIP-CR+) were comparable in the two nuclei (bottom).

### SST+ inhibitory cells

In the mouse BLA, SST is present in a significant number of GABAergic cells; however, their postsynaptic targets, morphological appearance and single-cell features are mostly unexplored. Therefore, we first examined the postsynaptic target distribution of SST- expressing axon terminals. We labeled SST-expressing inhibitory cells in Sst-Cre mice using a viral vector and then the sections containing the amygdala region were immunostained for a voltage-gated potassium channel Kv2.1, which visualizes the somata of amygdalar principal neurons (Vereczki et al., 2016). By counting the number of EYFP-expressing boutons that formed close appositions with the Kv2.1-immunoreactive somata, we observed that a very few of SST-expressing boutons target this membrane compartment of principal neurons either in the LA or BA (LA: 1 %, n = 667 boutons; BA: 1.8 %, n = 1388 boutons, n=2 mice). These results suggest that SST-containing boutons preferentially innervate the dendrites of neurons in mice similarly to that found in rats (Muller et al., 2007). To reveal the targets of SST- expressing axon endings and their occurrence along the dendritic tree of principal neurons, we intracellularly labeled single neurons in the LA and BA in acute slices that were prepared from mice where EYFP was expressed in SST-containing GABAergic cells. Using double immunostaining, we found that EYFP-expressing boutons that formed close appositions with the intracellularly labeled principal cells were evenly distributed along their dendritic trees (Figure 7A, B). LA (n =8) and BA (n = 9) principal cells were similarly covered by EYFP+ axon terminals (*p* = 0.28, Kolmogorov Smirnov test), therefore these results were pooled (Figure 7B). With a closer inspection, we observed that EYFP-expressing boutons overall targeted dendritic shaft more frequently than spines (Figure 7B). In LA, 137 and 814 boutons contacted spines and dendritic shaft, respectively (n = 8 principal cells, 12,176 µm total dendritic length), whereas 131 boutons formed close appositions with spines and 411 boutons with shafts of BA principal cells (n = 9, 13,816 µm total dendritic length). The ratio of boutons contacting shafts vs. spines in the LA (6.3 ± 1.1) and BA (4.3 ± 0.8) was similar (*p* = 0.16, Mann Whitney U test). The distribution of EYFP-expressing boutons targeting dendritic shaft or spines along the dendritic trees was not significantly different either (*p* = 0.16 in the LA, and *p* = 0.08 in the BA). Thus, our results confirmed that the vast majority of SST- containing axon terminals target the dendritic compartment of amygdalar principal cells (Muller et al., 2007; Wolff et al., 2014); consequently, these GABAergic cells are in a position to play a role in dendritic inhibition.

**Figure 7.**
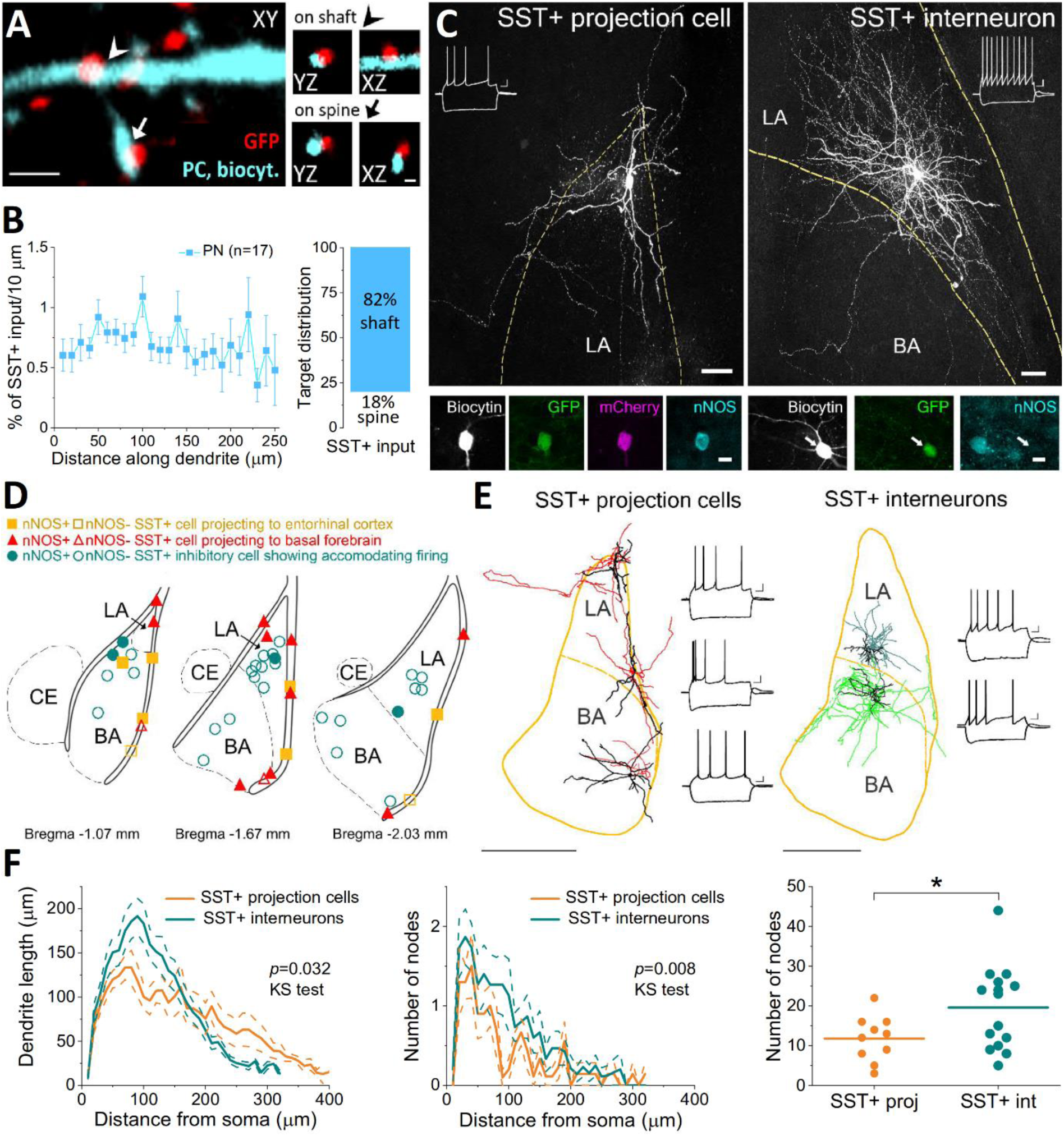
Characterization of GABAergic cells expressing SST in Sst-Cre mice. ***A,*** GFP+ boutons labeled in the LA and BA using a viral strategy in Sst-Cre mice form close appositions with a dendritic shaft (arrowhead) and spine (arrow) of an intracellularly labelled amygdalar principal neuron (PN)(scale bars = 4 and 1 µm for large and small images, respectively). ***B,*** *Left,* SST+ inputs evenly covered the dendritic tree of principal neurons (8 LA and 9 BA PNs). *Right*, The majority of SST+ boutons (n = 1493) targets dendritic shafts of amygdalar principal neurons. ***C,*** Two examples of biocytin-filled SST+ GABAergic cells recorded in acute slices. SST+ projection cells were labeled using an intersectional strategy by injecting retroAAV-mCherry-Flpo into the basal forebrain or entorhinal cortex and AAV- C(on)/F(on)-EYFF into the amygdala region. SST+ interneurons were visualized upon injection of AAV-DIO-EYFP into the BLA. Note that both SST+ projection cells and interneurons showed accommodation in their spiking and displayed a sag in their voltage responses upon negative step current injection (insets). In SST+ projection cells, nNOS was typically present, whereas SST+ interneurons were immunonegative for this enzyme (scale bars = 50 and 10 µm for large and small images, respectively). ***D,*** Distribution of SST+ inhibitory cells recorded in amygdalar slices. Each symbol represents the location of the cell body. ***E,*** Neurolucida reconstruction of three SST+ inhibitory cells projecting to the basal forebrain and two SST+ interneurons (dendrites in black, axons in color) and the corresponding voltage responses upon intracellular step current injections (scale bar = 500 µm). Note the elongated and less ramified dendrites of projection cells, and more branched axons of interneurons, and also the similar voltage responses of both types of SST+ inhibitory cells upon step current injections (x = 100 ms, y = 10 mV). ***F,*** Comparison of the structure of the dendritic trees of SST+ projection cells (n = 10) and interneurons (n = 15) using Sholl analysis. Dendritic length (left) and the number of nodes (center) as a function of distance from the soma are significant different. Mean, solid line; S.E.M., dashed line. Whereas the total dendritic length is comparable for the two SST+ inhibitory cell types, the total number of nodes is significantly higher for SST+ interneurons (*p* = 0.043). Number of nodes for individual neurons are represented by dots, while the lines indicate the mean.

In the next set of investigations, our goal was to characterize the GABAergic cells giving rise to SST-containing boutons in the two examined amygdalar nuclei. Previous studies showed that SST is expressed at least in two types of GABAergic cells. One type innervates primarily the dendrites of principal cells, whereas the other type, somata of which are present often in the external capsule, projects to remote areas, including the basal forebrain and entorhinal cortex (McDonald et al., 2012; McDonald and Zaric, 2015). Yet, none of these inhibitory cell types have been examined in details. To record from SST-expressing GABAergic projection cells, we applied intersectional strategy by injecting retroAAV-mCherry-Flpo viruses into the basal forebrain or entorhinal cortex of Sst-cre mice, and AAV-C(on)/F(on)-EYFP into the amygdala region. This approach revealed retrogradely labeled SST+ cells in the BLA (as well as in surrounding areas) in green, some of which were recorded and filled with biocytin in acute slices (Figure 7C). We sampled 28 and 16 GABAergic cells that projected to the basal forebrain or the entorhinal cortex, respectively. We included only those neurons in the analysis that had axon collaterals in the LA and/or BA (12 and 8 SST+ projection cells targeting the basal forebrain and entorhinal cortex, respectively), i.e. these GABAergic projection cells were in the position to participate in amygdala function. As neither the features of dendritic arbors, nor the single-cell characteristics were found to be different in inhibitory cells projecting to the basal forebrain or entorhinal cortex (*p* > 0.21), we pooled the two datasets. Previous findings indicated that SST+ GABAergic projection neurons often express nNOS (Sik et al., 1994; He et al., 2016), therefore, we tested the presence of this enzyme in the sampled neurons using immunostaining. Accordingly, the vast majority of SST+ GABAergic projection neurons indeed showed immunoreactivity for nNOS (78%, n = 18 tested). In both groups of projection neurons, 2-2 immunonegative cells were found. These observations therefore imply that the presence of nNOS in SST+ inhibitory cells may be a good tool to separate projection cells from those that are local interneurons.

Next, we aimed to compare the properties of these SST+ projection cells with SST+ interneurons. To this end, we randomly sampled green cells in acute slices that were prepared from the amygdala region of Sst-cre mice after viral labeling (Figure 7C). Total of 31 EYFP+ neurons were recorded with sufficiently labeled dendritic and/or axonal arbors (17 in the LA and 14 in the BA). Out of these green neurons, two were fast spiking basket cells (one of them showed immunopositivity for PV, whereas in the other case, the PV immunoreactivity could not be unequivocally determined) and one neurogliaform cell. These three neurons were excluded from further analyses. Using immunostaining, we tested the expression of nNOS in remaining EYFP+ neurons and found that this enzyme was present only in a minority of randomly sampled SST+ inhibitory cells (15%, n = 26; 2 in the LA and 2 in the BA, Figure 7D). As our results show that nNOS is often present in SST+ GABAergic projection cells, we excluded these 4 SST+/nNOS+ inhibitory cells from further comparisons, since they might have been randomly sampled projection neurons. Thus, the restricted group of SST+ interneurons was composed of 13 and 11 cells in the LA and BA, respectively. As the single cell features of SST+ interneurons located in the LA and BA were similar (*p* > 0.2), the results were pooled and compared to those obtained for SST+ GABAergic projection cells. We found that all, but one parameters investigated were similar in the two types of SST+ inhibitory cells (Figure 7C, E, Table 8).

**Table 8.** Single-cell properties of SST-expressing GABAergic cell types in the LA and BA. Data are presented as the median with the first and third quartiles in parentheses. P values obtained by Mann-Whitney (MW) U test are indicated. Significant difference is shown in bold.

| Parameters | SST+ interneurons<br>(n=17-24) | SST+ GABAergic proj. cells<br>(n=15-20) | MW test<br>p |
| --- | --- | --- | --- |
| AP half-width (ms) | 0.7<br>(0.6, 0.8) | 0.8<br>(0.7, 0.8) | 0.16 |
| Spike rate (Hz) | 45<br>(37.5, 51.3) | 55<br>(32.5, 72.5) | 0.37 |
| AHP 50% decay (ms) | 34.6<br>(9.2, 65.8) | 27.6<br>(13.5, 42.7) | 0.72 |
| ISI between the first two spikes (ms) | 9.2<br>(7.9, 11.85) | 7.6<br>(6.35, 10.3) | 0.17 |
| ISI between the last two spikes (ms) | 28.2<br>(21.55, 33.35) | 26.75<br>(16.3, 34.1) | 0.65 |
| Accommodation ratio | 2.98<br>(1.91, 4.29) | 2.86<br>(2.16, 4.48) | 0.93 |
| Input resistance (MOhm) | 252.8<br>(219.9, 362.6) | 279.3<br>(196.6, 388.3) | 0.97 |
| Membrane time constant (ms) | 24.75<br>(20.81, 33.4) | 20.8<br>(15.23, 25.7) | 0.053 |
| Membrane capacitance (pF) | 93.3<br>(76.7, 121.2) | 71.5<br>(55.5, 93.8) | **0.014** |
| Relative sag amplitude | 0.28<br>(0.134, 0.427) | 0.177<br>(0.125, 0.399) | 0.4 |
| Sag delay (ms) | 84.6<br>(62.3, 99.5) | 62.8<br>(49.4, 99.4) | 0.45 |

AP, action potential; AHP, after hyperpolarization, ISI, inter-spike interval.

During the inspection of intracellularly labeled SST+ inhibitory cells, we noticed that projection cells often emitted elongated dendrites and had only a few axon collaterals in slices, while local interneurons had rather multipolar dendritic trees and dense axonal arborization (Figure 7C, E). As the dendritic tree may be less impacted by slicing than axons, we compared only the features of dendrites in the two groups of SST+ inhibitory cells. We found no difference in any parameters for SST+ interneurons in the LA (n = 8) and BA (n= 7; *p* > 0.2), therefore the two datasets were pooled and compared to those obtained for SST+ projection cells (n = 10). Although no difference was found in the total dendritic length between SST+ projection cells (2, 467 ± 434 µm, n = 10) and interneurons (2,695 ± 234 µm, n = 15, *p* = 0.62), the structure of their dendritic trees was clearly different. Sholl analysis revealed that the dendritic branches of SST+ projection neurons were longer and less ramified (Figure 7F), which was also reflected in the total number of nodes (11.8 ± 1.8, n = 10 SST+ projection cells; 19.6 ± 2.7, n = 15 SST+ interneurons; *p* = 0.043). These results show that the two groups of SST+ inhibitory cells display distinct morphology.

In the last set of experiments, we attempted to estimate the ratio of SST+ GABAergic cells by labeling them in Sst-Cre mice using a viral vector (Figure 8A-E, Table 6). We found that the fraction of SST+ inhibitory cells differed significantly in the two amygdalar nuclei (Table 6). As two interneurons with a fast spiking phenotype were found among randomly sampled SST+ inhibitory cells, we checked the colocalization of PV and SST in the population of GABAergic cells: we found a negligible presence of this Ca^2+^ binding protein in SST+ inhibitory cells (LA: 1.3 %, n = 77, BA: 1.6 %, n = 185, n = 3 mice). Finally, we assessed the ratio of SST+ interneurons and SST+ GABAergic cells with long-range projections in virus-labeled cells in Sst-Cre mice using immunostaining against nNOS (Figure 8D). We observed that a considerable fraction of GFP-expressing SST+ neurons showed immunolabeling for nNOS+ (LA: 41.6 %. BA: 25 %), a ratio that was significantly different in the LA and BA (*p* = 0.043, Figure 8E, Table 6). These data indicate, in line with observations obtained in other cortical areas, that dendrite-targeting SST+ interneurons are more abundant than GABAergic projection cells expressing SST.

**Figure 8.**
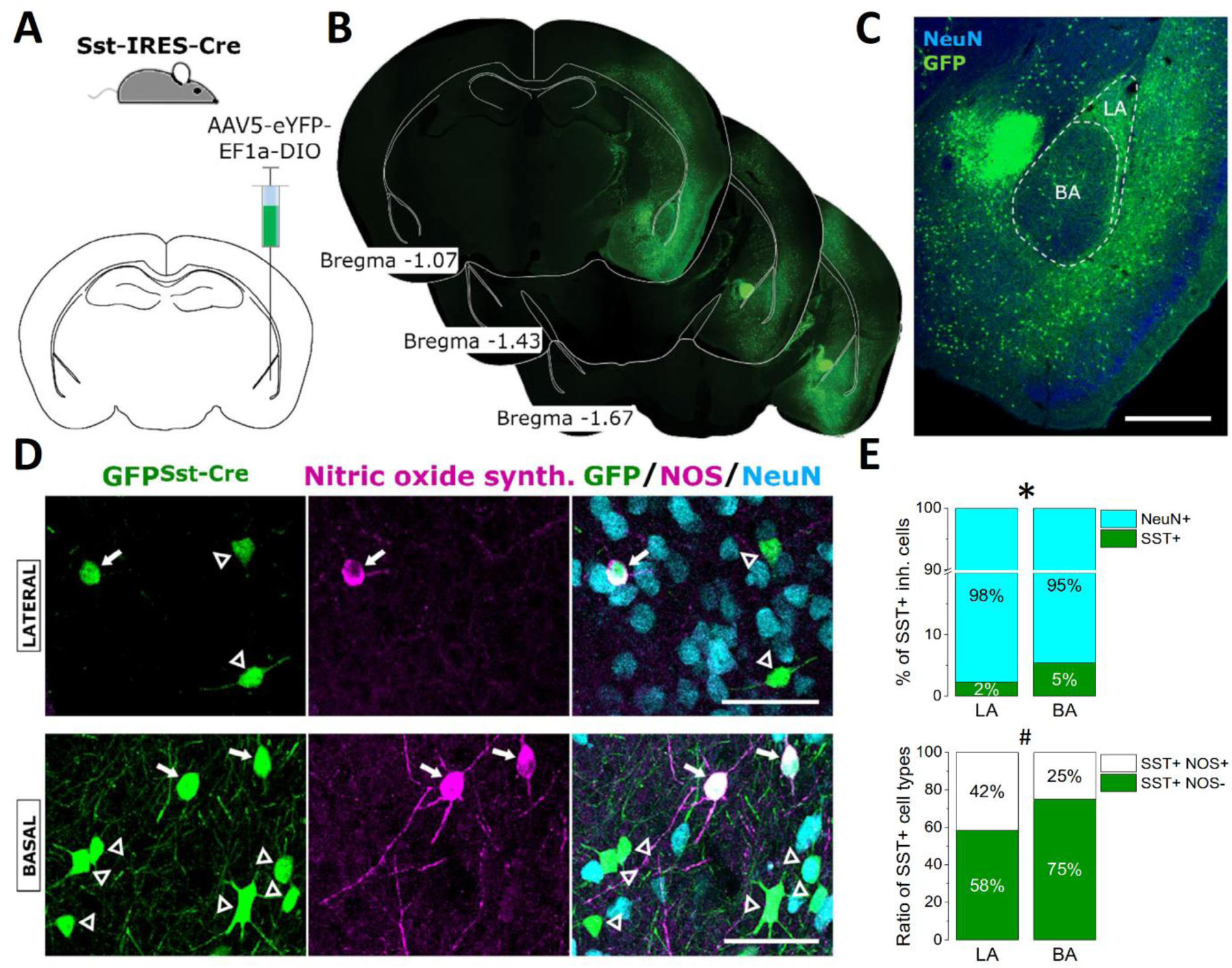
SST-expressing GABAergic cells in the LA and BA. ***A***, Schematic of the strategy for targeting SST-expressing GABAergic cells in the amygdala. ***B***, Representative images of GFP expression after virus transfection taken at the corresponding anterior-posterior coordinates (in mm) relative to Bregma. ***C***, Representative example of the amygdalar region taken at a higher magnification (scale bar = 500 µm). ***D***, In the majority of GFP-labeled neurons in Sst-Cre mice, no immunoreactivity for neuronal nitric oxide synthase (NOS, open arrowheads) was observed, yet there was a number of virus-labeled neurons that showed immunopositivity for this enzyme (arrows) in both nuclei (scale bars = 50 µm). ***E***, Both the percentage of SST+ GABAergic cells (top) and the percentage of SST+ inhibitory cells that express NOS (bottom) were different between the two nuclei (*, *p* = 0.0018; #, *p* =0.043).

### NPY+ neurogliaform cells

In the subsequent investigation, we aimed to estimate the fraction of neurogliaform cells in the two amygdalar nuclei. As NPY has been shown to be a characteristic marker for the vast majority, if not for all neurogliaform cells in cortical regions (Armstrong et al., 2012; Manko et al., 2012; Paul et al., 2017), we used Npy-Cre mice to label these GABAergic neurons in the amygdala by injecting viruses carrying EYFP. We found that the vast majority of labeled neurons in Npy-Cre mice had indeed morphological features typical for GABAergic neurons, but there were some labeled neurons with clear principal cell appearance. In line with this later notion, there was a considerable axonal projection in the contralateral BA in unilaterally injected Npy-Cre mice, axon collaterals that were decorated with boutons immunoreactive for VGluT1 (data not shown), a type of vesicular glutamate transporter expressed in amygdalar principal cells (Andrasi et al., 2017). In addition, principal cells could be recorded, although infrequently, in offspring of Npy-Cre x Ai14 mice (see below).

Thus, to ensure that we study only NPY+ GABAergic neurons in the amygdala, double-transgenic mice were generated by intercrossing Npy-Cre mice with Dlx5/6-Flpe mice that express Flp recombinase in the majority of GABAergic neurons in cortical structures (Miyoshi and Fishell, 2011). Then, we injected INTRSECT viruses into the amygdalar region of Npy^Cre^;Dlx5/6^Flp^ mice to transfect those GABAergic neurons with EYFP content that express both Cre and Flp recombinases. This approach resulted in no labeling in principal cells assessed by the lack of axonal projection in the contralateral BA and by sampling no principal cells in acute slice preparations.

To reveal the cell types that express NPY in the LA and BA, we performed whole-cell recordings in EYFP-expressing neurons in acute amygdalar slices that were prepared from AAV-injected Npy^Cre^;Dlx5/6^Flp^ mice or offspring of Npy-Cre x Ai14 mice. Based on the single-cell electrophysiological features, including the action potential half-width (measured at half the amplitude between the threshold and the peak voltage), maximum spiking rate, accommodation ratio, membrane time constant and sag amplitude, three main GABAergic cell groups could be identified among randomly sampled neurons expressing reporter proteins (Figure 9). The largest number of recorded neurons (52%) were neurogliaform cells showing a typical late-spiking phenotype (Figure 9A, D). The single-cell properties of these neurons could be characterized by wide action potentials, moderate spike rate with accommodation, fast membrane time constant and no or minimal sag in their voltage responses upon negative current injections (Figure 9E, Table 9). Labeled interneurons in this group displayed characteristic morphological features of neurogliaform cells, including short smooth dendrites that often ramified and dense local axonal arborization (Figure 9A). Many of these interneurons showed weak immunoreactivity for nNOS (22/25 tested), but none for PV (0/9 tested), CB (0/25 tested) or SST (0/4 tested)(Figure 9A). The second largest group of interneurons (33%) showed a fast spiking phenotype (Figure 9B, D). These interneurons had narrow spikes, the highest firing rate, no accommodation in spiking, fast membrane time constant and no sag (Figure 9E, Table 9). Spine-free dendrites and axon arborization of the interneurons in this category resembled the appearance typical for PV+ basket cells and axo-axonic cells (Vereczki et al., 2016). Immunostaining revealed that many of these NPY+ interneurons were indeed immunoreactive for PV (5/7 tested) and for CB (3/6 tested)(Figure 9B). Out of 6 fast spiking interneurons tested, we observed SST immunoreactivity in one case. In addition, where only a part of the axon could be revealed, labeled boutons of this fast spiking NPY+ interneuron formed close appositions with axon initial segments visualized by Ankyrin G staining, confirming that some of the axo-axonic cells can express NPY. The third group of recorded neurons (15%) had relatively narrow spikes, showed accommodation in spiking, had a relatively slow membrane time constant and a sag in their voltage responses upon negative current injections (Figure 9E, Table 9). Morphological characteristics of these GABAergic cells were similar to those typical for SST+ interneurons, including sparsely spiny dendrites and elongated soma. Indeed, all but one of the NPY+ GABAergic cells in this group showed strong immunopositivity for SST (6/7 tested)(Figure 9C). We have also recorded from an NPY+ interneuron, which showed clear immunoreactivity for CB1 on its axon terminals and displayed a typical firing pattern of CCK+ basket cells. In addition to inhibitory cells, some principal cells were also recorded in offspring of Npy-Cre x Ai14 mice (n=3), but not in AAV- injected Npy^Cre^;Dlx5/6^Flp^ mice. These results obtained in acute slices combined with immunocytochemical data suggest that the ratio of neurogliaform cells, PV+ fast spiking interneurons and SST+ inhibitory cells visualized in Npy^Cre^;Dlx5/6^Flp^ mice by a viral vector can be assessed by their PV or SST content at the population level.

**Figure 9.**
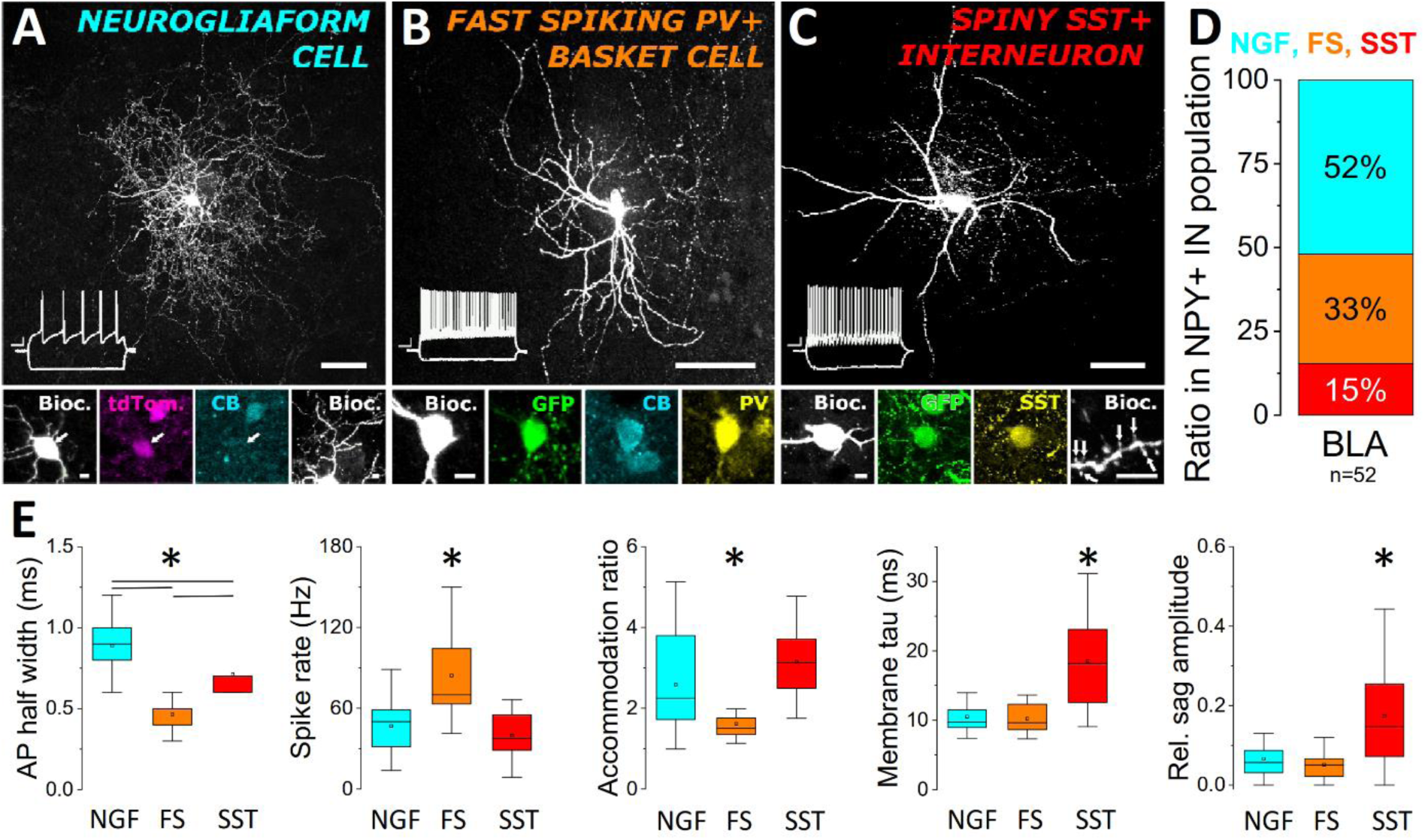
Three distinct inhibitory cell types express neuropeptide Y (NPY) in the amygdala. ***A,*** In amygdalar slices prepared from offspring of virus-injected Npy^Cre^;Dlx5/6^Flp^ mice or Npy-Cre x Ai 14 mice, the majority of recorded neurons had neurogliaform cell (NGF) morphology (a dense local axon arborization; short, frequently ramified dendrites (see small images below), a late-spiking phenotype (inset) and lacked calbindin (CB) content. ***B***, Another large group of NPY+ interneurons showed a fast spiking phenotype (inset) and was immunoreactive for parvalbumin (PV), and often for CB, which is typical for PV+ basket cells. ***C***, Inhibitory cells in the smallest group of NPY+ neurons had sparsely spiny dendrites (white arrows point to spines in the small image) and showed immunopositivity for somatostatin (SST). Firing of these interneurons showed accommodation and sag in their voltage responses upon negative step current injection (inset). Scale bars = 50 µm and 5 µm for large and small images, respectively. Scale bars of the firing patterns are x = 100 ms, y = 10 mV. ***D***, Ratio of the morphologically, neurochemically and electrophysiologically different NPY+ inhibitory cell types sampled *in vitro*. BLA here refers to LA and BA. ***E***, Single-cell properties of the three distinct GABAergic cells types expressing NPY in the amygdala. * indicates a significant difference, for data, see Table 9.

**Table 9.** Single-cell properties of the three NPY-expressing GABAergic cell types in the LA and BA. Data are presented as the median with the first and third quartiles in parentheses. P values obtained by statistical tests are indicated. Significant differences shown in bold and italic were determined by Kruskal-Wallis ANOVA and Mann-Whitney (MW) U test, respectively. NGF, neurogliaform cell; Fast spiking, fast spiking PV+ interneurons; SST+, SST-immunoreactive interneurons.

| Parameters | NGF<br>(n=26-28) | Fast spiking<br>(n=14-17) | SST+ (n=8) | Kruskal-<br>Wallis<br>ANOVA | NGF vs<br>FS MW<br>test | NGF vs<br>SST+<br>MW<br>test | FS vs<br>SST+<br>MW<br>test |
| --- | --- | --- | --- | --- | --- | --- | --- |
| AP half-width<br>(ms) | 0.9<br>(0.8, 1) | 0.5<br>(0.4, 0.5) | 0.7<br>(0.6, 0.7) | **<0.001** | *<0.001* | *0.007* | *<0.001* |
| Spike rate (Hz) | 50<br>(31.3, 58.8) | 70<br>(61.6, 114.1) | 37.55<br>(27.5, 58.75) | **<0.001** | *<0.001* | 0.38 | *0.0029* |
| AHP 50% decay<br>(ms) | 58.8<br>(38.7, 114.7) | 14.4<br>(10.1, 29.5) | 24.1<br>(9.62, 75.3) | **<0.001** | *<0.001* | 0.09 | 0.37 |
| ISI between the<br>first two spikes<br>(ms) | 9.65<br>(5.57, 18.17) | 8.2<br>(5.8, 11.17) | 9.8<br>(5.35, 13.37) | 0.56 |  |  |  |
| ISI between the<br>last two spikes<br>(ms) | 25.25<br>(19.9, 34.3) | 15.3<br>(5.9, 17.8) | 29<br>(17.62, 39.65) | **<0.001** | *<0.001* | 0.97 | *0.009* |
| Accommodation<br>ratio | 2.25<br>(1.72, 3.8) | 1.51<br>(1.35, 1.76) | 3.125<br>(2.42, 3.81) | **<0.001** | *0.003* | 0.16 | *<0.001* |
| Input resistance<br>(MΩ) | 177.55<br>(154.45, 234.7) | 133.4<br>(109.1, 205.5) | 233.3<br>(136.6, 302.8) | **0.033** | *0.011* | 0.57 | 0.11 |
| Membrane time<br>constant (ms) | 9.71<br>(8.89, 11.54) | 9.64<br>(8.52, 12.34) | 18.18<br>(12.52, 24.6) | **0.004** | 0.81 | *0.002* | *0.0037* |
| Membrane<br>capacitance (pF) | 53<br>(43, 62.7) | 68.4<br>(58.3, 84.8) | 76.7<br>(56.5, 137.1) | **0.004** | *0.007* | 0.014 | 0.56 |
| Relative sag<br>amplitude | 0.123<br>(0.096, 0.168) | 0.1<br>(0.068, 0.14) | 0.196<br>(0.112, 0.235) | **0.046** | 0.44 | *0.029* | *0.029* |

AP, action potential; AHP, after hyperpolarization, ISI, inter-spike interval.

Before performing this estimation, we first determined the fraction of all NPY+ inhibitory cells in the LA and BA (Figure 10, Table 6). The ratio of NPY+ inhibitory cells was significantly different in the two amygdalar nuclei (Table 6). To reveal the ratio of NPY+ inhibitory cells that express PV or SST, we tested the neurochemical content of GFP+ cells using immunostaining (Figure 10D, F). We observed that a significant fraction (∼ 25 %) of these neurons was immunoreactive for PV (Figure 10D, E, Table 6). In addition, a similarly large portion (∼ 27 %) of EYFP-expressing GABAergic neurons showed immunoreactivity against SST (Figure 10F, G, Table 6). Thus, based on the immunostaining, about half of the NPY+ GABAergic neurons (∼ 45% in both amygdalar nuclei) was immunoreactive neither for PV nor for SST, a group of neurons that should correspond to neurogliaform cells. Although there is almost no overlap between PV and SST immunoreactivity in amygdalar inhibitory cells, it would be more accurate to estimate the ratio of NPY+ neurons lacking PV and SST immunoreactivity in the same immunostained sections. Moreover, CB, which was absent in *in vitro* labeled neurogliaform cells, but is present in a large number of various types of GABAergic cells in the BLA (McDonald and Mascagni, 2001), may visualize additional inhibitory cell types, refining the ratio of neurogliaform cells even more. Therefore, we performed immunostaining against PV, SST and CB in amygdalar sections of Npy^Cre^;Dlx5/6^Flp^ mice in which EYFP expression visualized GABAergic cells. Using this approach, we found that neurogliaform cells may represent appr. 45 % of NPY+ GABAergic cells expressing EYFP but lacking immunoreactivity for PV, SST and CB (4 sections/mouse, 3 mice)(Figure 10G, Table 6). Thus, based on the observations that almost half of all NPY+ GABAergic cells are neurogliaform cells, we calculated that these interneurons make up 1.8 % and 3.5 % of all neurons in the LA and BA, respectively.

**Figure 10.**
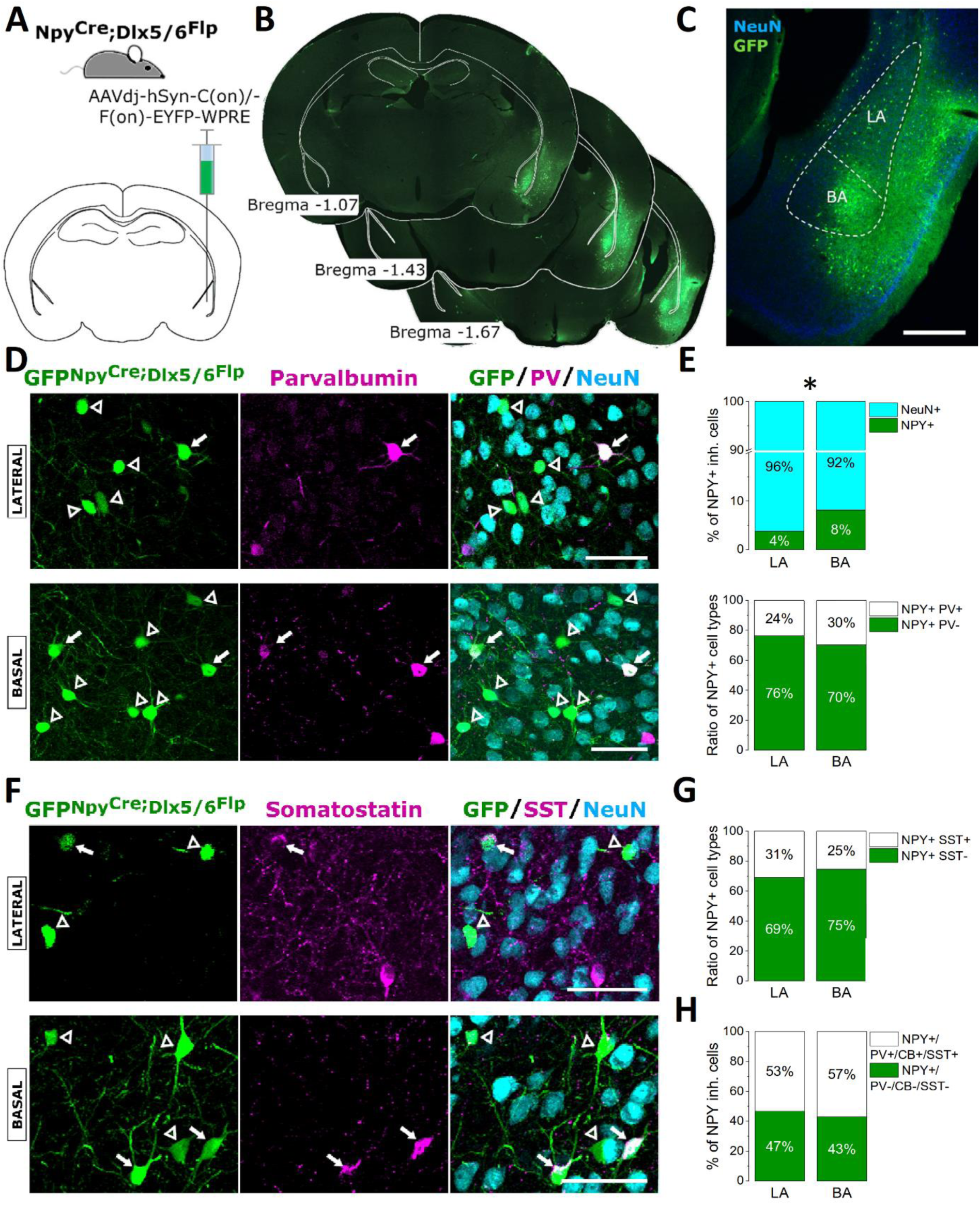
Neuropeptide Y (NPY)-expressing GABAergic cells in the LA and BA. ***A***, Schematic of the strategy for targeting NPY-expressing interneurons in the amygdala. ***B***, Representative images of GFP expression after virus transfection taken at the corresponding anterior-posterior coordinates (in mm) relative to Bregma. ***C***, Representative example of the amygdalar region taken at a higher magnification (scale bar = 500 µm). ***D***, In Npy^Cre^;Dlx5/6^Flp^ mice, a notable portion of GFP-labeled GABAergic cells expresses parvalbumin (PV)(arrows) in both nuclei. Open arrowheads indicate GFP+ neurons lacking PV immunoreactivity (scale bar = 50 µm). ***E***, The ratio of NPY+ GABAergic cells in the LA and BA was significantly different (top, *, *p* < 0.001), but the ratio of NPY+ inhibitory cells containing PV did not differ (bottom). ***F***, Another portion of NPY+ GABAergic cells showed immunoreactivity for somatostatin (SST)(arrows) in both nuclei. Open arrowheads indicate GFP+ neurons lacking SST immunoreactivity (scale bar = 50 µm). ***G***, The proportion of NPY+ and SST+ inhibitory cells was comparable in the LA and BA. ***H***, Almost half of the virus-labeled neurons in Npy^Cre^;Dlx5/6^Flp^ mice should represent the population of neurogliaform cells both in the LA and BA, assessed by lack of immunoreactivity in GFP-labeled interneurons for PV, calbindin (CB) and SST.

**Figure 11.**
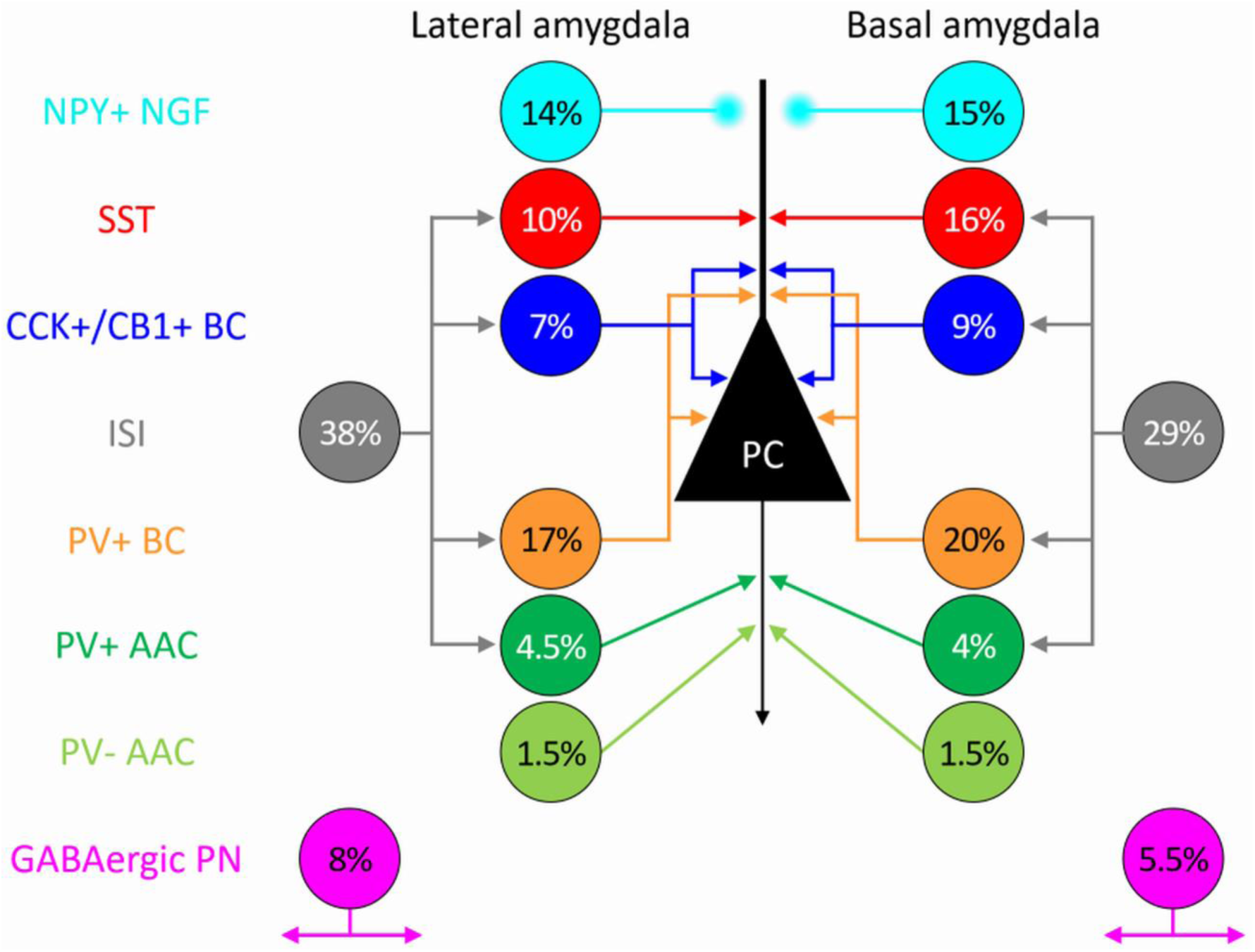
Ratios of the major inhibitory neuron types within the investigated GABAergic cell population of the LA and BA. NPY+ NGF, NPY-expressing neurogliaform cells; SST, somatostatin-expressing dendrite-targeting interneurons; CCK+/CB1+ BC, cholecystokinin and CB1 cannabinoid receptor type 1-expressing basket cells; ISI, interneuron-selective interneurons; PV+ BC, parvalbumin-containing basket cells; PV+ AAC, parvalbumin-expressing axo-axonic cells; PV-AAC, parvalbumin-lacking axo-axonic cells; GABAergic PN, GABAergic projection neurons expressing somatostatin; PC, principal cell. Based on previously published data, connectivity of inhibitory cells is shown. Arrows indicate the classical synaptic contacts, whereas circle for neurogliaform cell connections refers to the loose contacts typically formed by these GABAergic cells.

### The ratios of the major inhibitory cell categories are similar in the LA and BA

Finally, we calculated the fractions of the major GABAergic cell types within the investigated inhibitory cell groups in the two amygdalar nuclei. Our results show that these cell types constitute a comparable fraction of inhibitory neurons in the LA and BA (Figure 11).

## Discussion

In the mouse, we found that the number of neurons in the LA is significantly less than that found in the BA, in agreement with a recent study using molecular biological approaches (O’Leary et al., 2020). This observation is in contrast with findings reported in the rat, monkey and human amygdala, where a similar or rather larger number of neurons were observed in the LA compared to the BA (Schumann and Amaral, 2005; Chareyron et al., 2011). The discrepancy between the ratios of neurons in the LA and BA reported in the mouse in comparison to previous studies investigating other mammals may reflect diversity across species. Alternatively, defining the BA borders may substantially differ among studies, which may explain the differences at least in part. In other studies, amygdalar nuclei including the BA were usually defined based on cytoarchitecture, whereas here we used VAchT immunostaining to objectively delineate the BA borders, an approach that may be easy to adopt in future studies.

The ratio of inhibitory cells in the LA and BA found in this study is similar to those estimated earlier in rat and monkey BLA (McDonald, 1992; McDonald and Augustine, 1993) and are in good agreement with the overall estimation of GABAergic cell number in other cortical structure (Gabbott and Somogyi, 1986; Beaulieu et al., 1992; Ren et al., 1992). At present, it is unclear the reason why the LA contains substantially less inhibitory cells, but one may speculate that this difference in neuronal composition supports distinct roles for the LA and BA in amygdala-related circuit operations (Janak and Tye, 2015; Manassero et al., 2018).

In addition to the number of GABAergic cells, we also attempted to assess the ratio of distinct inhibitory neuron types in the LA and BA. We took advantage of the fact that in adult Pvalb-Cre, Sst-Cre and Vip-Cre mice viral labeling visualizes one or two GABAergic cell categories that can be separated by immunostaining. In Pvalb-Cre mice, fast spiking basket cells and axo-axonic cells could be distinguished based on the CB content (Bienvenu et al., 2012; Vereczki et al., 2016; Andrasi et al., 2017; Rovira-Esteban et al., 2019). We estimated that 17 % and 20 % of all GABAergic cells are PV+ basket cells in the LA and BA, respectively. Similar ratio for this interneuron type was estimated in the hippocampus (Bezaire and Soltesz, 2013), whereas twice as many PV-expressing interneurons were found in the frontal, primary somatosensory and visual cortices (Xu et al., 2010). The large difference in the ratio in interneurons containing PV suggest that the convergence and divergence between these interneurons and their targets may follow distinct rules in different cortical areas.

One of our novel findings is the identification of axo-axonic cells lacking PV in the LA and BA. Importantly, axo-axonic cells expressing or lacking PV followed the same innervation strategy in both nuclei, confirming and expanding our earlier observations (Veres et al., 2014). Based on our data, it is safe to predict that PV is absent in ca.1/3 of all axo-axonic cells in the LA and BA. This ratio posits these two amygdalar nuclei between the hippocampus and prefrontal cortex, as in the former structure only PV-containing axo-axonic cells have been found (Katsumaru et al., 1988), whereas in the latter the majority of these GABAergic interneurons lacks PV (Wang et al., 2019).

In this study, we used a novel transgenic mouse line, the BAC-CCK-GFPcoIN_sb to visualize CCK-expressing GABAergic cells in the brain after intercrossing with Vgat-Cre. In offspring (i.e. Vgat^Cre^;CCK-GFPcoIN mice), the majority of recorded interneurons were basket cells. We found that 7-9 % of all inhibitory cells in the LA and BA are CCK+/CB1+ basket cells, a ratio, which is similar to that estimated in the hippocampus (Bezaire and Soltesz, 2013). Of note, in this novel transgenic mouse line we have not sampled fast spiking cells or neurogliaform cells, interneuron types that could be often targeted in Cck^Cre^;Dlx5/6^Flp^ mice, in addition to CCK+/CB1+ basket cells (Rovira-Esteban et al., 2019). These data suggest that the two strategies, the knock-in *vs.* the use of BAC as a tool to generate transgenic lines, produce mice that express the Cre recombinase or fluorescent proteins in distinct populations of CCK+ inhibitory cells, yet CB1+ basket cells are always affected, albeit with a different efficacy.

In Vip-Cre mice, interneuron-selective interneurons containing or lacking CR can be predominantly, if not exclusively labeled using the method applied (Rhomberg et al., 2018; Krabbe et al., 2019). Based on earlier data obtained in the hippocampus (Acsády et al., 1996; Gulyas et al., 1996; Hájos et al., 1996), we hypothesized that CR+/VIP- interneurons also belong to the interneuron-selective interneuron group, although future work should confirm our assumption. One of our surprising findings was that interneuron-selective interneurons in the LA (38 %) and BA (29 %) are more abundant than in the hippocampus (20 %)(Bezaire and Soltesz, 2013) and neocortex (23-30 %)(Xu et al., 2010). Thus, our observation may imply that the massive regulatory potential of interneuron-selective interneurons over other GABAergic cells can play a central role in the control of various amygdala functions, as it has been shown recently for affective memory formation (Krabbe et al., 2019).

In this study, we provide the first detailed characterization of SST+ GABAergic cells in the LA and BA. As in the hippocampus and neocortex (Katona et al., 1999; Wang et al., 2004), SST+ inhibitory cells target predominantly the dendritic shaft and to a lesser extent, the spines of principal cells. SST+ GABAergic cells that project to the basal forebrain or entorhinal cortex (McDonald et al., 2012; McDonald and Zaric, 2015) were found to be immunopositive for nNOS. This enzyme content in SST+ GABAergic cells, thus, helped us to estimate the ratios of interneurons and projection neurons expressing SST in the LA and BA. We found a similar ratio for SST+ GABAergic projection cells in the amygdala as it was reported in the hippocampus (5-6 %)(Bezaire and Soltesz, 2013). However, this latter study estimated significantly less SST+ interneurons in the hippocampus (4-5%), than we found in the amygdala (10-16 %), or others in the neocortex (17-20 %)(Xu et al., 2010). Future studies should clarify the reason of this surprisingly low ratio of SST+ interneurons in the hippocampus.

NPY has been shown to be expressed often in neurogliaform cells (Fuentealba et al., 2010; Tricoire et al., 2010; Armstrong et al., 2012; Manko et al., 2012; Perrenoud et al., 2013). In neocortical areas, 7-10 % of GABAergic cells was found to express NPY (Xu et al., 2010), whereas around 30 % of all inhibitory cells may belong to neurogliaform cell family in the hippocampus (Bezaire and Soltesz, 2013). Thus, the LA and BA, where we estimated 14-15 % of GABAergic cells to be neurogliaform cells, take up an intermediate position between these two cortical structures. Our observation that Cre recombinase under the control of NPY is expressed in a portion of PV+ basket and axo-axonic cells in the two amygdalar nuclei examined is novel, but not surprising, as in the hippocampus NPY immunoreactivity has been reported in some PV+ interneurons (Klausberger et al., 2004), whereas many SST+ GABAergic cells express NPY in cortical regions (He et al., 2016; Lim et al., 2018).

Adding up the fractions of each GABAergic cell type resulted in a sum, which is close to the ratios of GABAergic cells obtained by unbiased stereological analysis. This notion strongly suggests that the vast majority of GABAergic cells in the LA and BA belong to the seven cardinal inhibitory cell categories examined in this study. In addition to these GABAergic cells typical for cortical structures, other inhibitory cell types do exist in the BLA, like those expressing muscarinic acetylcholine receptor type 2 (M2), but they do not provide a large number of GABAergic cells (McDonald and Mascagni, 2011).

Our present study in mice determined the number of GABAergic and non-GABAergic neurons in the LA and BA as well as provided a realistic estimate for the proportions of distinct inhibitory cell types. These results will pave the ground for future studies, specifically for those aiming to reveal the changes in amygdalar inhibitory circuits in different models of neuropsychiatric diseases, including anxiety, autism spectrum disorder and schizophrenia. The significance of these investigations are highlighted by the fact that hyper-excitability in the amygdala, arising from the imbalance between excitation and inhibition typifies many pathological brain states in humans (Rosen and Schulkin, 1998; Rosenkranz et al., 2010; Prager et al., 2016; Sharp, 2017; Takarae and Sweeney, 2017).

## Acknowledgements

We acknowledge financial support from the Intramural Program of Semmelweis University awarded to VKV, from the National Research, Development and Innovation Office (K_119742) awarded to NH and from the Hungarian Brain Research Program (2017-1.2.1-NKP-2017-00002) awarded to NH. The authors are grateful to Erzsébet Gregori for her excellent technical assistance. We also thank László Barna, the Nikon Microscopy Center at the Institute of Experimental Medicine, Nikon Austria GmbH, and Auro-Science Consulting, Ltd., for kindly providing microscopy support.

## Author contributions

Designed experiments: NH, VKV

Performed experiments: VKV, KM, EK, ZM, ZF, LRE

Analyzed data: VKV, KM, EK, ZF, LRE, JMV, HN

Supervised the project: VKV, EF, NH

Wrote the paper: NH

